# Genome-wide locus sequence typing (GLST) of eukaryotic pathogens

**DOI:** 10.1101/2020.03.24.003590

**Authors:** Philipp Schwabl, Jalil Maiguashca Sánchez, Jaime A. Costales, Sofía Ocaña, Maikell Segovia, Hernán J. Carrasco, Carolina Hernández, Juan David Ramírez, Michael D. Lewis, Mario J. Grijalva, Martin S. Llewellyn

## Abstract

Analysis of genetic polymorphism is a powerful tool for epidemiological surveillance and research. Powerful inference from pathogen genetic variation, however, is often restrained by limited access to representative target DNA, especially in the study of obligate parasitic species for which *ex vivo* culture is resource-intensive or bias-prone. Modern sequence capture methods enable pathogen genetic variation to be analyzed directly from vector/host material but are often too complex and expensive for resource-poor settings where infectious diseases prevail. This study proposes a simple, cost-effective ‘genome-wide locus sequence typing’ (GLST) tool based on massive parallel amplification of information hotspots throughout the target pathogen genome. The multiplexed polymerase chain reaction amplifies hundreds of different, user-defined genetic targets in a single reaction tube, and subsequent agarose gel-based clean-up and barcoding completes library preparation at under 4 USD per sample. Approximately 100 libraries can be sequenced together in one Illumina MiSeq run. Our study generates a flexible GLST primer panel design workflow for *Trypanosoma cruzi*, the parasitic agent of Chagas disease. We successfully apply our 203-target GLST panel to direct, culture-free metagenomic extracts from triatomine vectors containing a minimum of 3.69 pg/μl *T. cruzi* DNA and further elaborate on method performance by sequencing GLST libraries from *T. cruzi* reference clones representing discrete typing units (DTUs) TcI, TcIII, TcIV, and TcVI. The 780 SNP sites we identify in the sample set repeatably distinguish parasites infecting sympatric vectors and detect correlations between genetic and geographic distances at regional (< 150 km) as well as continental scales. The markers also clearly separate DTUs. We discuss the advantages, limitations and prospects of our method across a spectrum of epidemiological research.

## Introduction

Genome-wide single nucleotide polymorphism (SNP) analysis is a powerful and increasingly common approach in the study and surveillance of infectious disease. Understanding patterns of SNP diversity within pathogen genomes and across pathogen populations can resolve fundamental biological questions (e.g., reproductive mechanisms in *T. cruzi*^1^, reconstruct past^2^ and present transmission networks (e.g., *Staphylococcus* infections within hospitals)^3^ or identify the genetic bases of virulence^4,5^ and resistance to drugs (see examples from *Plasmodium* spp.^6,7^). A number of obstacles, however, complicate access to representative, genome-wide SNP information using modern sequencing tools. Micro-pathogens are often sampled in low quantities and together with large amounts of host/vector tissue, microbiota, or environmental DNA. Sequencing is rarely viable directly from the infection source and studies have often found it necessary to isolate and culture the target organism to higher densities before extracting DNA. These additional steps, however, are resource-intensive and bias-prone. Pathogen isolation is less often attempted on asymptomatic infections and is less likely to succeed when levels of parasitaemia in a sample are low. Genomic sequencing data on the protozoan parasite *Leishmania infantum*, for example, has for such reasons come to exhibit major selection bias towards aggressive strains isolated by invasive sampling from canine hosts. A short look into the limited number of whole-genome sequencing (WGS) datasets available for *L. infantum* at the European Nucleotide Archive (ENA) quickly confirms this statement. Vector-isolated genomes have yet to be reported from the Americas and only a single study claims to have sequenced *L. infantum* from asymptomatic hosts^8^. Selection bias also often occurs due to competition among isolated strains. Studies on the kinetoplastid *Trypanosoma cruzi*, for example, are time and again confounded by growth and survival rate differences among genotypes in culture^9–11^, and gradual reductions to genetic diversity are often observed over time^12^. Karyotypic changes are also known to arise during *T. cruzi* micromanipulation and axenic growth^13,14^.

A variety of approaches therefore aim to obtain genome-wide SNP information without first performing pathogen isolation and culturing steps. Some studies separate target sequences from total DNA or RNA by exploiting base modifications or transcriptional properties specific to the pathogen^15^, vector^16^ or host^17,18^. Others describe the use of biotinylated hybridization probes^19–22^ or selective whole-genome amplification, e.g., based on the strand displacement function of phi29 DNA polymerase^23^. Such techniques are costly and often excessive when a study’s primary objective is to evaluate genetic distances and diversity among samples rather than to reconstruct complete haplotypes or investigate structural genetic traits. Epidemiological tracking and source attribution studies, for example, often benefit little from measuring invariant sequence areas or defining the complete architecture of sample genomes. Also pathogen typing or population assignment objectives primarily require information on polymorphic sites. It is nevertheless quite common to see such studies to undertake expensive WGS procedures only for final analyses to take place ‘post-VCF’^24^, i.e., using a list of diagnostic markers compiled from a small fraction of polymorphic reads.

Highly multiplexed polymerase chain reaction (PCR) amplicon sequencing offers a much more efficient option when obtaining genome-wide SNP information is the primary goal. First marketed under the name Ion AmpliSeq by Thermo Fisher Scientific^25^, the method consists in the simultaneous amplification of dozens to hundreds of DNA targets known or hypothesized to contain sequence polymorphism in the sample set. Each sample’s resultant amplicon pool is then prepared for sequencing by index/adaptor ligation or in a subsequent ‘barcoding’ PCR. Panel construction is highly flexible, requiring only that the primers exhibit similar melting/annealing temperatures and a low propensity to cross-react. As such, target selection can be tailored to specific research goals, for example, to profile resistance markers^26^ or to genotype neutral SNP variation for landscape genetic techniques^27^. The potential to isolate and genotype pathogen DNA at high-resolution directly from uncultured sample types by multiplexed amplicon sequencing has however received little attention thus far. Simultaneous PCR-based detection of multiple pathogen species or genotypes is certainly common^28^, but multiplexable primer panels are rarely designed for subsequent sequencing and polymorphism analysis. The Ion AmpliSeq brand currently offers pre-designed panels for studies on ebola^29^ and tuberculosis^30^ but the use of custom panels for other pathogen species (e.g., *Bifidobacterium*^31^ or human papilloma virus^32^) remains surprisingly rare in the literature.

In this study we describe the design and implementation of a large multiplexable primer panel for *T. cruzi*, parasitic agent of Chagas disease. In contrast to past multi-locus sequence typing (MLST) methods involving at most 32 (individually amplified) gene fragments, our ‘genome-wide locus typing’ (GLST) tool simultaneously amplifies 203 sequence targets across 33 (of 47) *T. cruzi* chromosomes. We apply GLST to metagenomic DNA extracts from triatomine vectors collected in Colombia, Venezuela and Ecuador and further describe method sensitivity/specificity by sequencing GLST libraries from *T. cruzi* clones representing discrete typing units (DTUs) TcI, TcIII, TcIV, and TcVI. The 780 SNP sites identified from GLST amplicon sequencing repeatably distinguish parasites infecting sympatric vectors and detect correlations between genetic and geographic distances at regional (< 150 km) and continental scales. The markers also clearly separate DTUs. We discuss the advantages and limitations of our method for epidemiological studies in resource-poor settings where Chagas and other ‘neglected tropical diseases’ prevail.

## Methods

### Triatomine samples and *T. cruzi* reference clones

*T. cruzi*-infected intestinal tract and/or faeces samples of *Rhodnius ecuadoriensis* and *Panstrongylus chinai* were collected by the Centro de Investigación para la Salud en América Latina (CISeAL) in Loja Province, Ecuador, following protocols described in Grijalva et al. 2012^33^. DNeasy Blood and Tissue Kit (Qiagen) was used to extract metagenomic DNA. Infected intestinal material of *Panstrongylus geniculatus*, *R. pallescens* and *R. prolixus* from northern Colombia was also collected in previous projects^34–36^, likewise using DNeasy Blood and Tissue Kit to extract metagenomic DNA. *Panstrongylus geniculatus* specimens from Caracas, Venezuela were collected by the citizen science triatomine collection program (http://www.chipo.chagas.ucv.ve/vista/index.php) at Universidad Central de Venezuela. This program has supported various epidemiological studies in the capital district^37–39^. DNA was extracted from the insect faeces by isopropanol precipitation. Geographic coordinates and ecotypes (domestic, peri-domestic, or sylvatic) of the sequenced samples are provided in Supplementary Tbl. 1.

*T. cruzi* epimastigote DNA from reference clones Chile c22 (TcI) Arma18 cl. 1 (TcIII), Saimiri3 cl. 8 (TcIV), Para7 cl. 3 (TcVI), Chaco9 col. 15 (TcVI) and CL Brener (TcVI) was obtained from the London School of Hygiene & Tropical Medicine (LSHTM). DNA extractions at LSHTM followed Messenger et al. 2015^40^.

Uninfected *Rhodnius prolixus* gut tissue samples used for mock infections (see ‘Method development and library preparation’) were also provided by LSHTM. Special thanks to C. Whitehorn and M. Yeo for supervising dissections. Insects were euthanized with CO_2_ and hindguts drawn into 5 volumes of RNAlater (Sigma-Aldrich) by pulling the abdominal apex toward the posterior with sterile watchmaker’s forceps.

*T. cruzi* TcI X10/1 Sylvio reference clone (‘TcI-Sylvio’) epimastigotes used for mock infections and various other stages of method development were obtained from CISeAL. Cryo-preserved cells were returned to log-phase growth in liver infusion tryptose (LIT) and quantified by hemocytometer before pelleting at 25,000 g. Pellets were washed twice in PBS and parasites killed by resuspension in 10 volumes of RNAlater. DNA from these *T. cruzi* cells (and their dilutions with preserved *T. prolixus* intestinal tissue) was extracted by isopropanol precipitation.

Isopropanol precipitation was also used to extract DNA from *T. cruzi* plate clone TBM_2795_CL2. This sample was previously analyzed by WGS^1^ and served as a control for GLST method development in this study.

### GLST target and primer selection

We began our GLST sequence target selection process by screening single-nucleotide variants previously identified in *T. cruzi* populations from southern Ecuador^1^. Briefly, Schwabl et al. sequenced genomic DNA from 45 cloned and 14 non-cloned *T. cruzi* field isolates on the Illumina HiSeq 2500 platform and mapped resultant 125 nt reads to the TcI-Sylvio reference assembly using default settings in BWA-mem v0.7.3^41^. Single-nucleotide polymorphisms (SNPs) were summarized by population-based genotype and likelihood assignment in Genome Analysis Toolkit v3.7.0^42^, excluding sites with low cumulative call confidence (QUAL < 1,500) and/or aberrant read-depth (< 10 or > 100) as well as those belonging to clusters of three or more SNPs. A ‘virtual mappability’ mask^43^ was also applied to avoid SNP inference in areas of high sequence redundancy in the *T. cruzi* genome. Read-mapping and variant exclusion criteria were verified by subjecting TcI-Sylvio Illumina reads from Franzen et al. 2012^44^ to the same pipelines as the Ecuadorian dataset. An additional mask was set around small insertion-deletions suggested to occur in these reads based on the assumption that the reference sample should not present alternate genotypes in high-quality contigs of the assembled genome.

We extracted 160 nt segments from the *T. cruzi* reference genome (.fasta file) whose internal sequence (positions 41 to 120) contained between one and ten of 75,038 SNPs identified in the above WGS dataset. These 56,428 segments were further filtered for synteny between *T. cruzi* and *Leishmania major* genomes as defined by the OrthoMCL algorithm at TriTrypDB^45^. Such conserved segments may be least prone to repeat-driven nucleotide diversity and as such most amenable to PCR^46^. The 6,259 synteny segments found by OrthoMCL therefore proceeded to primer search with the high-throughput primer design engine BatchPrimer3^47^. As target SNPs did not occur in the outer 40 nt of each synteny segment, these flanking regions provided additional flexibility to identify primers matching the following criteria:

- min. size = 24 nt
- max. size = 35 nt
- optimal size = 24 nt
- min. product size = 120 nt
- max. product size = 160 nt
- optimal product size = 120 nt
- min. melting temperature = 63 °C,
- max. melting temperature = 65 °C,
- optimal melting temperature = 63 °C,
- max. self-complementarity: 4 nt
- max. 3’ self-complementarity: 2 nt
- max. length of mononucleotide repeats = 3 nt
- min. GC content = 40%
- max. GC content = 60%

Each of 286 forward primer candidates output by BatchPrimer3 received the additional 5’ tag sequence 5’-ACACTGACGACATGGTTCTACA-3’ and reverse primer candidates received the 5’ tag sequence 5’-TACGGTAGCAGAGACTTGGTCT-3’. These tag sequences enable single-end barcode and Illumina P5/P7 adaptor attachment in second-round PCR. Next, we determined binding energies (ΔG) for all possible primer-pairs using the primer compatibility software MultiPLX v2.1.4. We discarded primers with inter-quartile ranges crossing a threshold of ΔG = −12.0 kcal/mol. Primers with 20 or more interactions showing ΔG ≤ −12.0 kcal/mol were also disallowed. The remaining 248 primer-pairs (median ΔG = −9.0) underwent a last filtering step by screening for perfect matches in raw WGS sequence files (.fastq). Low match frequency led to the elimination of 45 additional primer pairs. WGS alignments corresponding to the 203 sequence regions targeted by this final primer set were visualized in Belvu v12.4.3^48^. The 403 SNPs occurring within these sequence regions distributed evenly across individuals in Loja Province. Using the ‘nj’ function from the ‘ape’ package v5.0 in R v3.4.1^49^, the 403 SNPs also reproduced neighbor-joining relationships observed based on total polymorphism identified by WGS (Supplementary Fig. 1). These observations lent further support to the suitability of the GLST marker panel for the analysis of genetic differentiation at the landscape-scale. The GLST sequence target selection process described above is summarized in Fig. 1.

**Figure 1.**
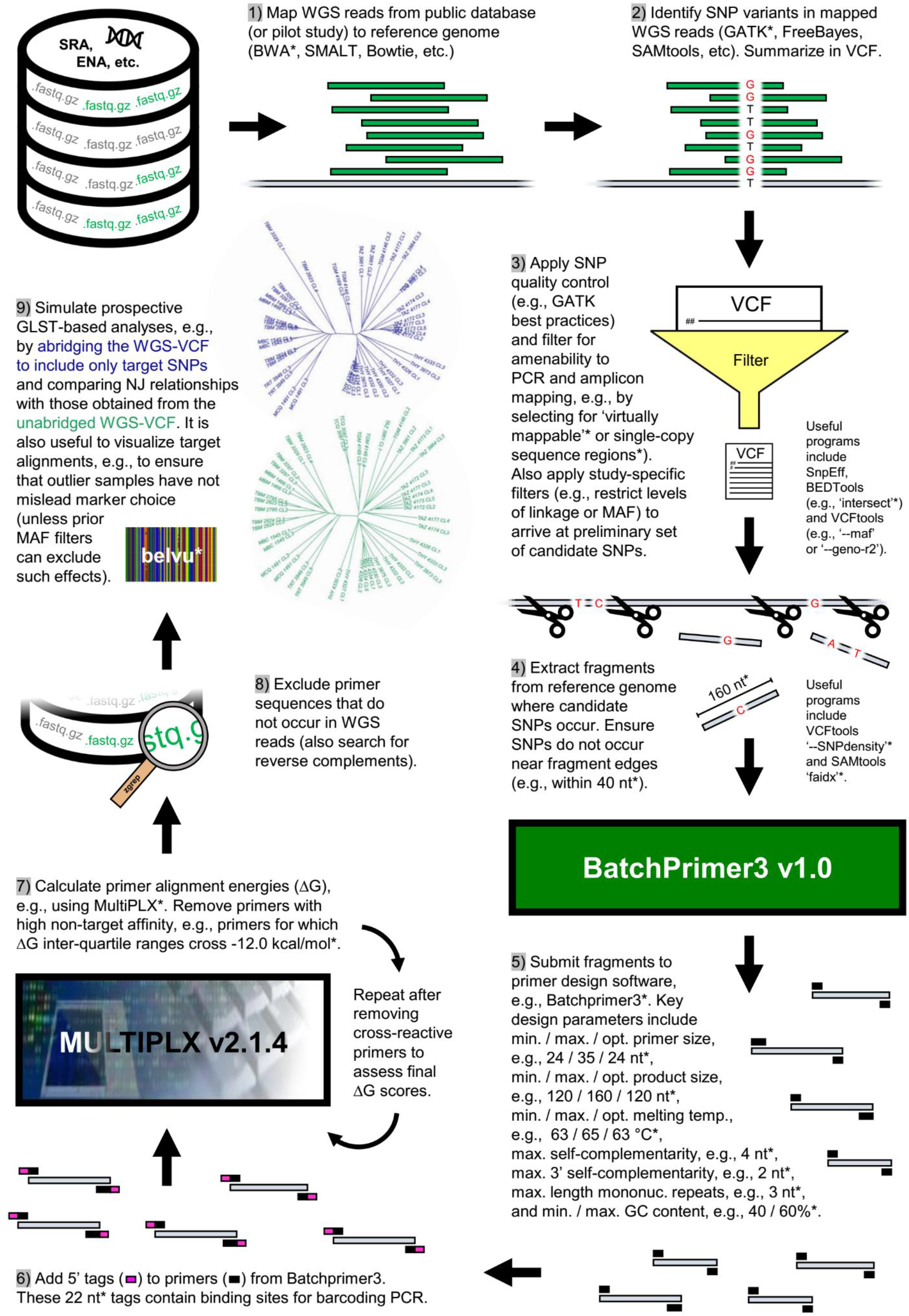
GLST sequence target selection from preliminary genomic data. Nine steps of primer panel construction and validation run clockwise from top left. Various methods and criteria can be applied to complete many of these steps. Those specific to this study are asterisked, e.g., we used BWA in step 1 and GATK in step 2. Abbreviations: SRA (Sequence Read Archive at www.ncbi.nlm.nih.gov/sra); ENA (European Nucleotide Database at www.ebi.ac.uk/ena; WGS (whole-genome sequencing); SNP (single-nucleotide polymorphism); MAF (minor allele frequency); PCR (polymerase chain reaction); VCF (variant call format); NJ (neighbor-joining).

### Wet lab method development and library preparation

The 203 primers pairs designed above (Supplementary Tbl. 2) were purchased from Eurofins Genomics (Ebersberg, Germany) at 200 μM concentration in salt-free, 96-well plate format. Primer pairs were first tested individually to establish cycling conditions for PCR (Supplementary Fig. 2). Optimal target amplification occurred with an initial incubation step at 98 °C (2 min); 30 amplification cycles at 98 °C (10 s), 60 °C (30 s), and 72 °C (45 s); and a final extension step at 72 °C (2 min). The 10 μl reactions contained 5 μl Q5 High-Fidelity Master Mix (New England Biolabs), 1 μl forward primer [10 μM], 1 μl reverse primer [10 μM], and 3 μl TcI-Sylvio epimastigote DNA. The multiplexed, first-round ‘GLST’ PCR reaction was prepared by combining all 406 primers in equal proportions and diluting the combined mix to 50.75 μM, resulting in individual primer concentrations of 50.75 μM / 406 = 125 nM. GLST reactions incorporated 2 μl of this primer mix rather than two separate 1 μl forward/reverse primer inputs as above.

We first tested GLST PCR on DNA extracts from mock infections, each consisting of 10^4^, 10^5^ or 10^6^ TcI-Sylvio epimastigote cells and one uninfected *R. prolixus* intestinal tract (Supplementary Fig. 3). Amplicons from lower concentration epimastigote dilutions gave weaker signals in gel electrophoresis, suggesting lower infection load thresholds at which vector gut DNA becomes unsuitable for GLST. Most vector gut DNA extracts obtained for this study represented donated material of limited quality and infection load, some samples were also without signal in PCR spot tests for the presence of high frequency ‘TcZ’^50^ satellite DNA (commonly targeted to diagnose human *T. cruzi* infections).

We therefore first used qPCR to identify vector gut samples containing *T. cruzi* DNA quantities within ranges successfully visualized from GLST reactions on epimastigote DNA quantified by Qubit fluorometry (Invitrogen) and serially diluted from 1.35 ng/μl to 2.50 pg/μl in dH_2_O (Supplementary Fig. 4). Each 20 μl qPCR reaction consisted of 10 μl SensiMix SYBR Low-ROX reagent (Bioline), 1 μl TcZ forward primer (5’-GCTCTTGCCCACAMGGGTGC-3’)^50^ [10 μM], 1 μl TcZ reverse primer (5’-CCAAGCAGCGGATAGTTCAGG-3’)^50^ [10 μM], 7 μl dH_2_O, and 1 μl vector gut DNA. Samples were amplified together with a 15-step standard curve containing between 0.30 pg and 4.82 ng *T. cruzi* epimastigote DNA. Reaction conditions consisted of an initial incubation step at 95 °C (10 min) and 40 amplification cycles at 95 °C (15 s), 55 °C (15 s), and 72 °C (15 s). Fluorescence acquisition occurred at the end of each cycle and final product dissociation was measured in 0.5 °C increments between 55 and 95 °C.

Vector gut samples suggested to contain at least 1.0 pg/μl *T. cruzi* concentrations based on qPCR proceeded to final library construction (Supplementary. Tbl. 1) alongside DNA from *T. cruzi* clones TBM_2795_cl2 (TcI), Chile c22 (TcI) Arma18 cl. 1 (TcIII), Saimiri3 cl. 8 (TcIV), Para7 cl. 3 (TcV), Chaco9 col. 15 (TcVI) and CL Brener (TcVI). Several samples were processed in 2 – 4 replicates beginning with the first-round GLST PCR reaction step. First-round PCR products were electrophoresed in 0.8% agarose gel to separate target bands (mode =164 nt) from primer polymers quantified with the Agilent Bioanalyzer 2100 System (see 78 nt primer peak in Supplementary Fig. 5). Excised target bands were resolubilized with the PureLink Quick Gel Extraction Kit (Invitrogen) to create input for subsequent barcoding PCR. This second PCR reaction consisted of an initial incubation step at 98 °C (2 min); 7 amplification cycles at 98 °C (30 s), 60 °C (30 s), and 72 °C (1 min); and a final extension step at 72 °C (3 min). Only 7 amplification cycles were used given polymer ‘daisy-chaining’ observed when cycling at 13 and 18x (Supplementary Fig. 6). The barcoding reaction adds Illumina flow cell and sequencing primer binding sites to each first-round PCR product. A different reverse primer is used for each sample. The reverse primer (5’-CAAGCAGAAGACGGCATACGAGAT*X*TACGGTAGCAGAGACTTGGTCT-3’) contains a 10 nt barcode (*X*) to distinguish reads from different samples during pooled sequencing. It also contains CS2 (sequencing primer binding sites). A single forward primer (5’-AATGATACGGCGACCACCGAGATCTACACTGACGACATGGTTCTA-3’) containing CS1 is used for all samples. Each 20 μl barcoding reaction contained 10 μl Q5 High-Fidelity Master Mix (New England Biolabs), 0.8 μl forward (universal) primer [10 μM], 0.8 μl (barcoded) reverse primer [10 μM], 5.4 μl dH_2_O and 3 μl (gel-purified) first-round PCR product. Barcoding primers were purchased from Eurofins Genomics at 100 μM concentration in HPLC-purified, 96-well plate format. Barcoded amplicons (e.g., Supplementary Fig. 7) were quantified by Qubit fluorometry (Thermo Fisher Scientific), and pooled at equimolar concentrations, gel-excised, re-solubilized, and verified by microfluidic electrophoresis (Supplementary Fig. 8) as above.

### GLST amplicon sequencing and variant discovery

The GLST pool was sequenced twice on an Illumina MiSeq instrument. We first used the pool to ‘spike’ additional base diversity into a collaborator’s 16S amplicon sequencing run. 16S samples were loaded to achieve 80% sequence output whereas GLST and PhiX DNA^51^ were each loaded at 10%. This first run occurred in 500-cycle format using MiSeq Reagent Kit v2. The second run occurred in 300-cycle format using MiSeq Reagent Micro Kit v2 and was dedicated solely to GLST (also no PhiX). Both runs were performed at Glasgow Polyomics using Fluidigm Custom Access Array sequencing primers FL1 (CS1 + CS2) and CS2rc^52^.

Demultiplexed sequence reads were trimmed to 120 nt and mapped to the TcI-Sylvio reference assembly using default settings in BWA-mem v0.7.3. Mapped reads with poor alignment scores (AS < 100) were discarded to decontaminate samples of non-*T.cruzi* sequences sharing barcodes with the GLST dataset. Identical results were achieved using BWA-sw in DeconSeq v0.4.3^53^ to decontaminate reads. After merging alignment (.bam) files from sequencing runs 1 and 2 with Picard Tools v1.11^54^, single-nucleotide polymorphisms (SNPs) were identified in each sample using the ‘HaplotypeCaller’ algorithm in GATK v3.7.0^42^. Population-based genotype and likelihood assignment followed using ‘GenotypeGVCFs’. We excluded SNP sites with QUAL < 80, D < 10, Mapping Quality (MQ) < 80 and or Fisher Strand Bias (FS) > 10. Individual genotypes were set to missing (./.) if they contained < 10 reads and set to reference (0/0) if they contained only a single alternate read (i.e., if they were classified as heterozygotes based on minor allele frequencies ≤ 10%). These filtering thresholds were cleared by all expected SNPs (i.e., SNPs also found in prior WGS sequencing) but not by all new SNPs found using GLST (e.g., see comparison of QUAL density curves in Supplementary Fig. 9). SNP calling with GATK was also performed separately for sequencing runs 1 and 2 in order to exclude SNP sites uncommon to both analyses from the merged dataset described above.

### GLST repeatability, population genetic and spatial analyses

We used PopART v1.7 to plot genetic differences between samples and sample replicates as a median-joining network, i.e., a minimum spanning tree composed of observed sequences and unobserved (reconstructed) sequence nodes^55^. Genetic differences were measured by applying the ‘vcf-to-tab’ script from VCFtools v0.1.13 to the filtered SNP dataset, concatenating each sample’s output fields and counting the number of mismatching alleles (0, 1 or 2) per site and sample pair. A phylogenetic tree was built by counting the number of non-reference alleles in each genotype with the VCFtools function ‘--012’, summing pairwise Euclidean distances at biallelic sites and plotting neighbor-joining relationships with the ‘nj’ function from the ‘ape’ package v5.0 in R v3.4.1^49^.

Considering only the first replicate of multiply sequenced samples, linkage and neutrality statistics were calculated using VCFtools functions ‘--geno-r2’ (calculates correlation coefficients between genotypes following Purcell et al.^56^), ‘--het’ (calculates inbreeding coefficients using a method of moments^57^) and ‘--hwe’ (filters sites by deviation from Hardy-Weinberg Equilibrium following Wigginton et al.^58^). F_ST_ differentiation was calculated using ARLSUMSTAT v3.5.2^59,60^.

Correlations between geographic and genetic differences were also calculated from non-reference allele counts in R v3.4.1^49^. The ‘mantel’ function from the ‘vegan’ package v2.4.4^61^ was used to test significance of the Mantel statistic by permuting geographic distances and re-measuring correlations to genetic distances 999 times. Again, we used only the first replicate for samples with replicate sets. DTU reference clones were also excluded from analysis. Geographic distances were measured by projecting sample latitude/longitude (WGS 84) coordinates into a common xy plane (EPSG code 3786) selected following Šavrič et al. 2016^62^ (Supplementary Tbl. 1). EPSG 3786 projection was also used to map samples with the Natural Earth quick start kit in QGIS v2.18.4.

Given that missing information in sequence alignment can confound inference on genetic distances between samples^63^, above repeatability and phylogenetic analyses excluded SNP sites in which genotypes were missing for any individual, and mantel analyses excluded SNP sites in which genotypes were missing in > 10% individuals. These exclusion criteria initially led to significant information loss due to the presence of two outlier samples, ARMA18_CL1_rep2 and COL253, libraries of which had been sequenced despite poor target visibility in gel electrophoresis (i.e., final PCR product banding appeared similar to that of ECU2 in Supplementary Fig. 7). Read-depths for the two samples ended up averaging 1.2 interquartile ranges below the sample set median and precluded genotype assignment at > 25% SNP sites. We therefore decided to exclude them from all analyses.

## Results

### SNP polymorphism and repeatability

GLST amplicons contained a total of 780 SNP sites, 387 polymorphic among TcI samples and 393 private to non-TcI reference clones (Fig. 2). Median read-depth was 266x across all sites. Of 403 loci targeted from the WGS dataset^1^, 97% (391) were recovered by GLST and 82 contained polymorphism outside of Ecuador. GLST recovered 80 of 87 SNPs previously identified in TBM_2795_CL2 using WGS. Minimum parasite DNA concentration successfully genotyped from metagenomic DNA was 3.69 pg/μl (sample ECU36 – see Supplementary Fig. 10).

**Figure 2.**
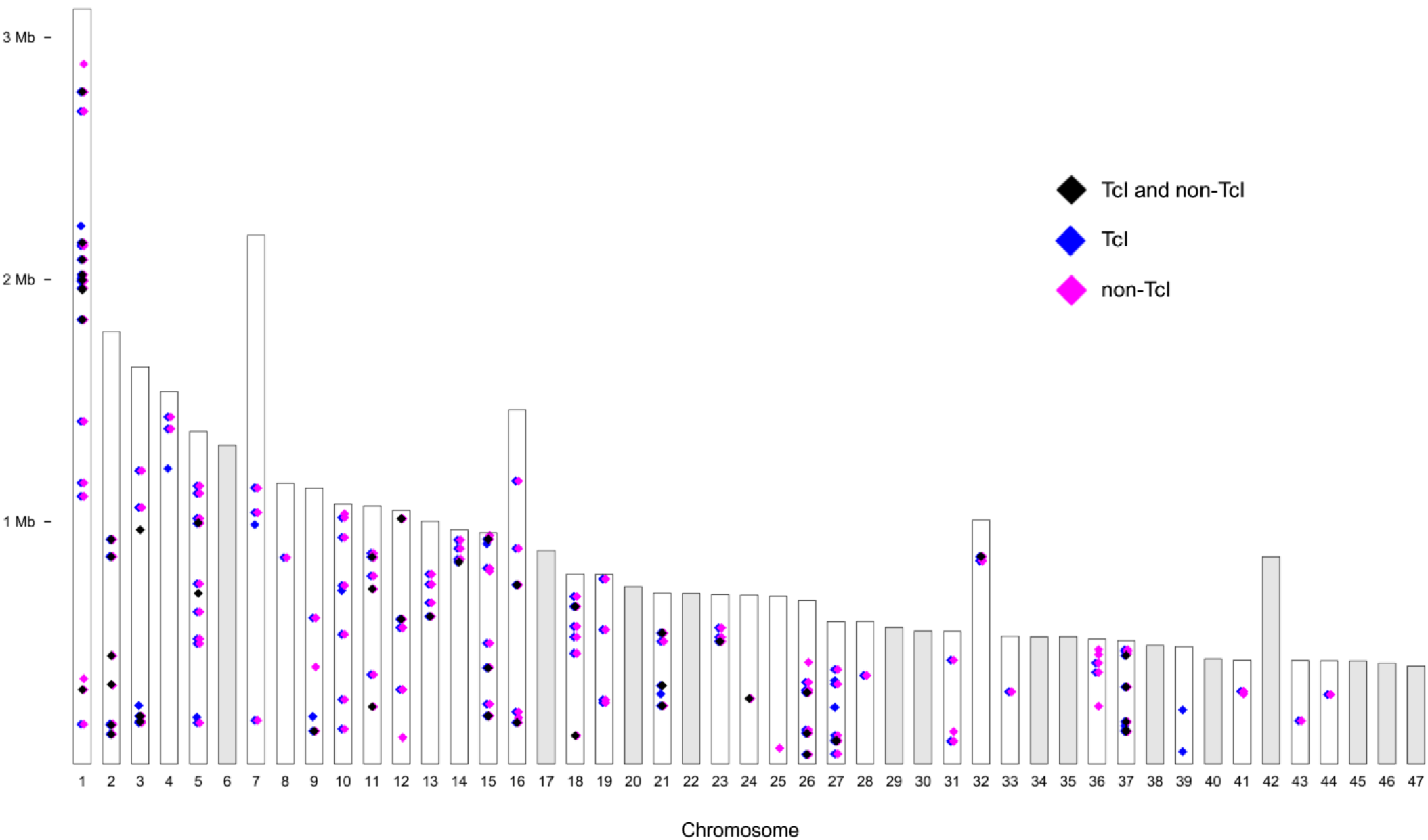
Variant loci detected in *T. cruzi* I samples and reference clones of other sub-lineages. The genome-wide distribution of SNP variants is shown relative to the TcI-Sylvio reference assembly. Each column represents one of 47 putative chromosomes. Pink diamonds comprise 393 variants that occur only in non-TcI samples. The remaining 387 variants are private to (blue) or shared by TcI and other sub-lineages (black). Diamonds representing nearby SNPs (e.g., those occurring on the same GLST target segment) overlap at this scale.

The TBM_2795_CL2 control sample underwent GLST in four replicates. These replicates were identical at all 561 SNP sites for which genotypes were called in all samples of the dataset. Median number of allelic differences (AD = 0, 1 or 2 per site) at non-missing sites between other replicate pairs was 3 (Tbl. 1). Pairwise AD did not correlate to minimum, maximum or difference in mean read-depth between the two replicates (p < 0.80).

Read-mapping coverage was inconsistent among replicates but strongly correlated between sequencing runs (Pearson’s r = 0.93, p < 0.001) (Supplementary Figs. 11 – 12). Variant calling was also highly consistent: prior to variant filtration, only 10 SNP sites were called from run1 that were not also called from run 2 (these were excluded from analysis – see Methods).

### Differentiation among *T. cruzi* individuals, sampling areas and sub-lineages

Sampling sites in Colombia, Venezuela and Ecuador are plotted in Fig. 3, and a median-joining network of allelic differences among GLST genotypes is shown in Fig. 4. GLST clearly distinguished TcI individuals at common collection sites in Soata (COL466 vs. COL468, AD = 37), Paz de Ariporo (COL133 vs. COL135, AD = 33), Tamara (COL154 vs. COL155 AD = 107) and Lebrija (COL77 vs. COL78, AD = 43) municipalities of Colombia but not in the community of Bramaderos (ECU3 vs. ECU8 vs. ECU10, AD = 0) in Loja Province, Ecuador. Samples from nearby sites within Caracas, Venezuela were also clearly distinguished by GLST (e.g., VZ16816 vs. VZ17114, AD = 43).

**Figure 3.**
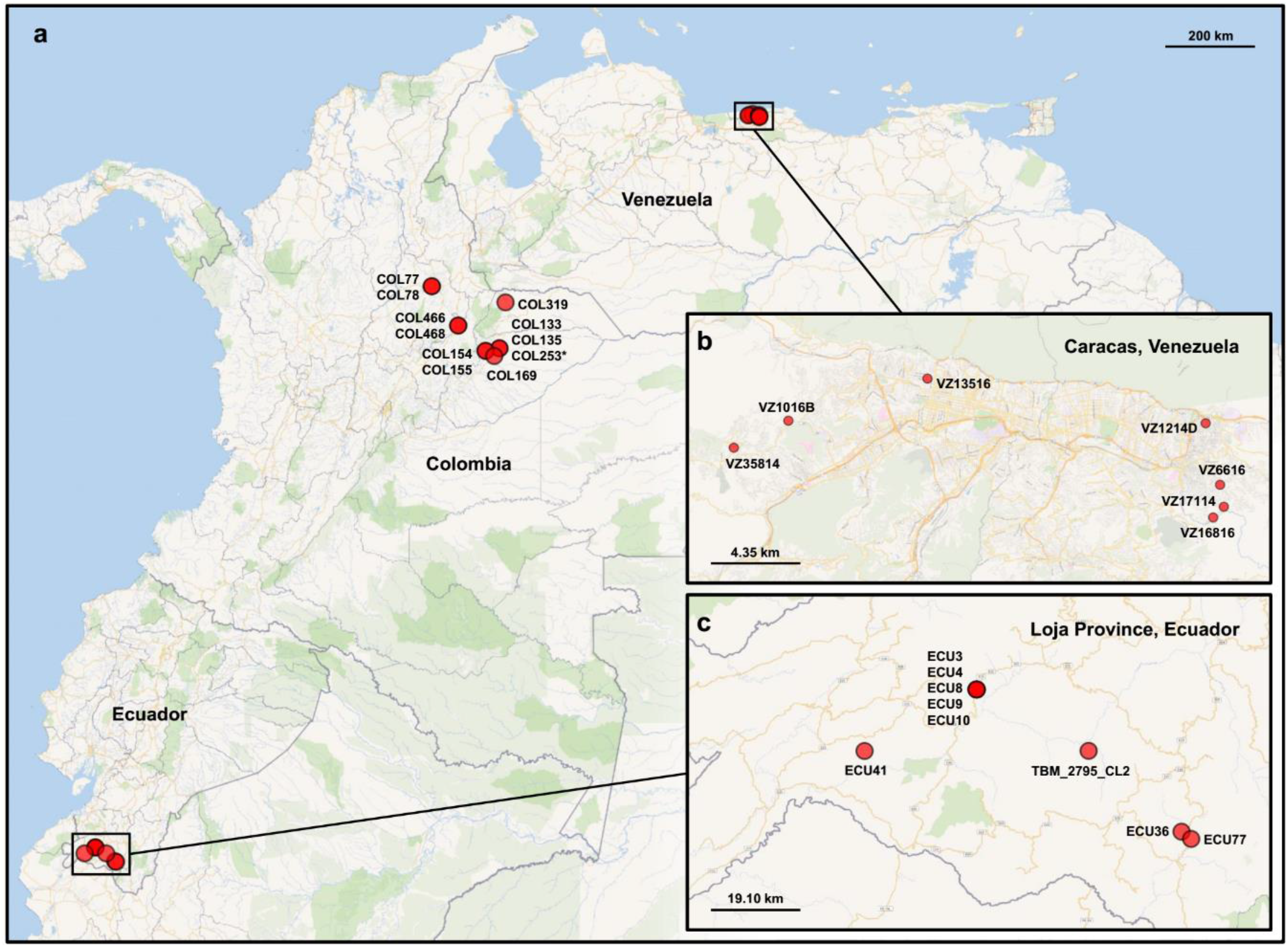
Map of vector sampling sites. **a** Sampling in Colombia involved a larger spatial area than that in Venezuela and Ecuador. *T. cruzi*-infected intestinal material was collected from *Panstrongylus* and *Rhodnius* vectors in Arauca, Casanare, Santander and Boyacá. We asterisk COL253 because low read-depth led to sample exclusion. **b***P. geniculatus* material from Venezuela was collected within the Metropolitan District of Caracas. **c** *R. ecuadoriensis* and *P. chinai* material from Ecuador was collected in Loja Province. Supplementary Tbl. 1 lists coordinates and other details.

**Figure 4.**
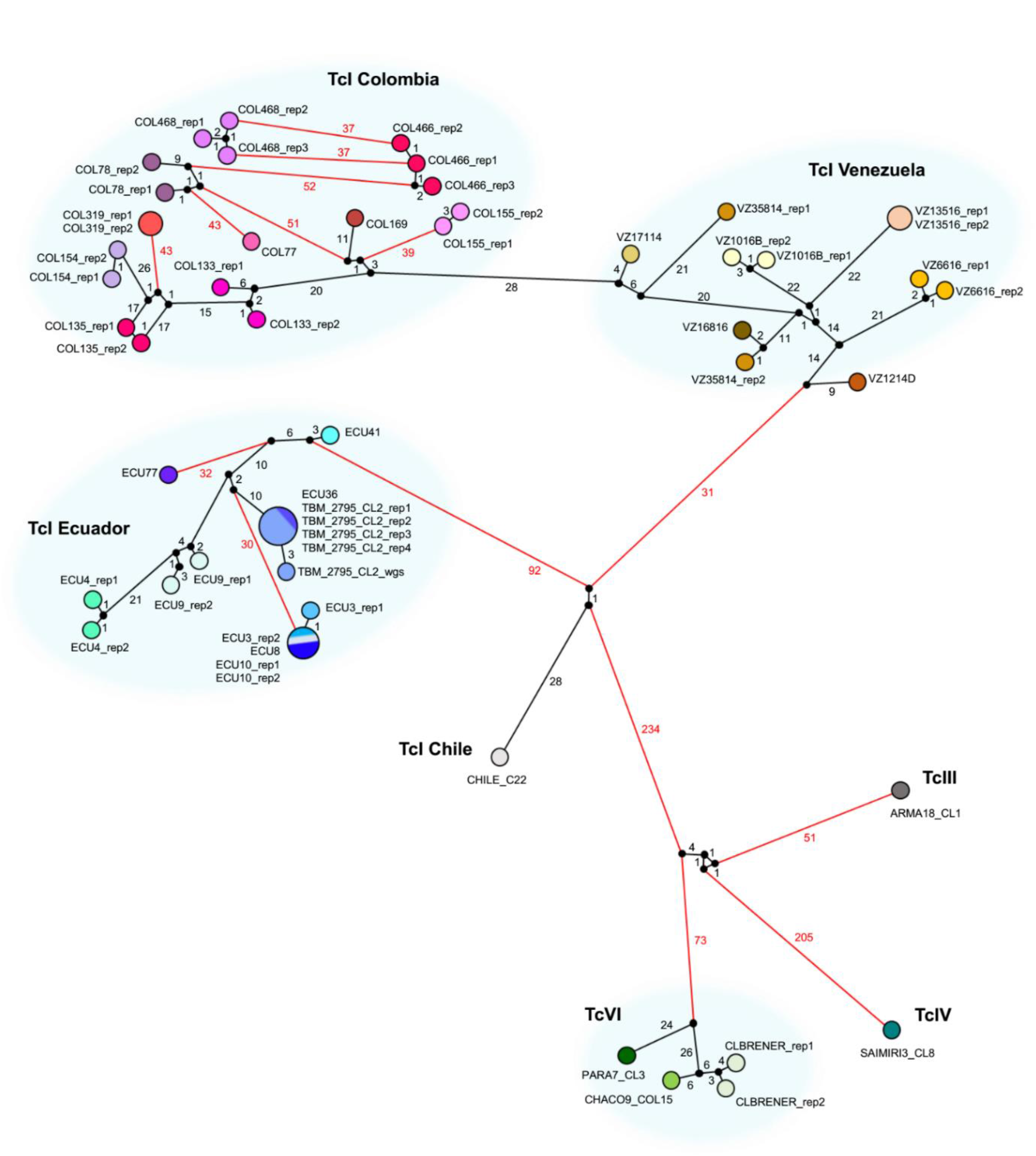
Allelic differences among *T. cruzi* I samples and reference clones of other sub-lineages as a median-joining network. A single SNP locus can differ by 0, 1 or 2 between two individuals (i.e., the individuals match at both, one, or neither allele). The AD measurement indicated on each edge of the network represents the total number of differences across all loci for which genotypes were called in all individuals of the dataset (n = 561). Red edges indicate differences of 30 and above. Technical replicates are represented by circles of the same fill color. Larger circles represent the occurrence of identical GLST genotypes. Edge length is not directly proportional to AD.

Nucleotide diversity (π = mean pairwise AD) was higher in samples from Caracas (π = 29.0) than in those from Loja Province (π = 22.8) but not in those from Colombia (π = 43.2) (Tbl. 2). Hardy-Weinberg ratios, linkage and inbreeding coefficients are also listed in Tbl. 2.

**Table 1.**
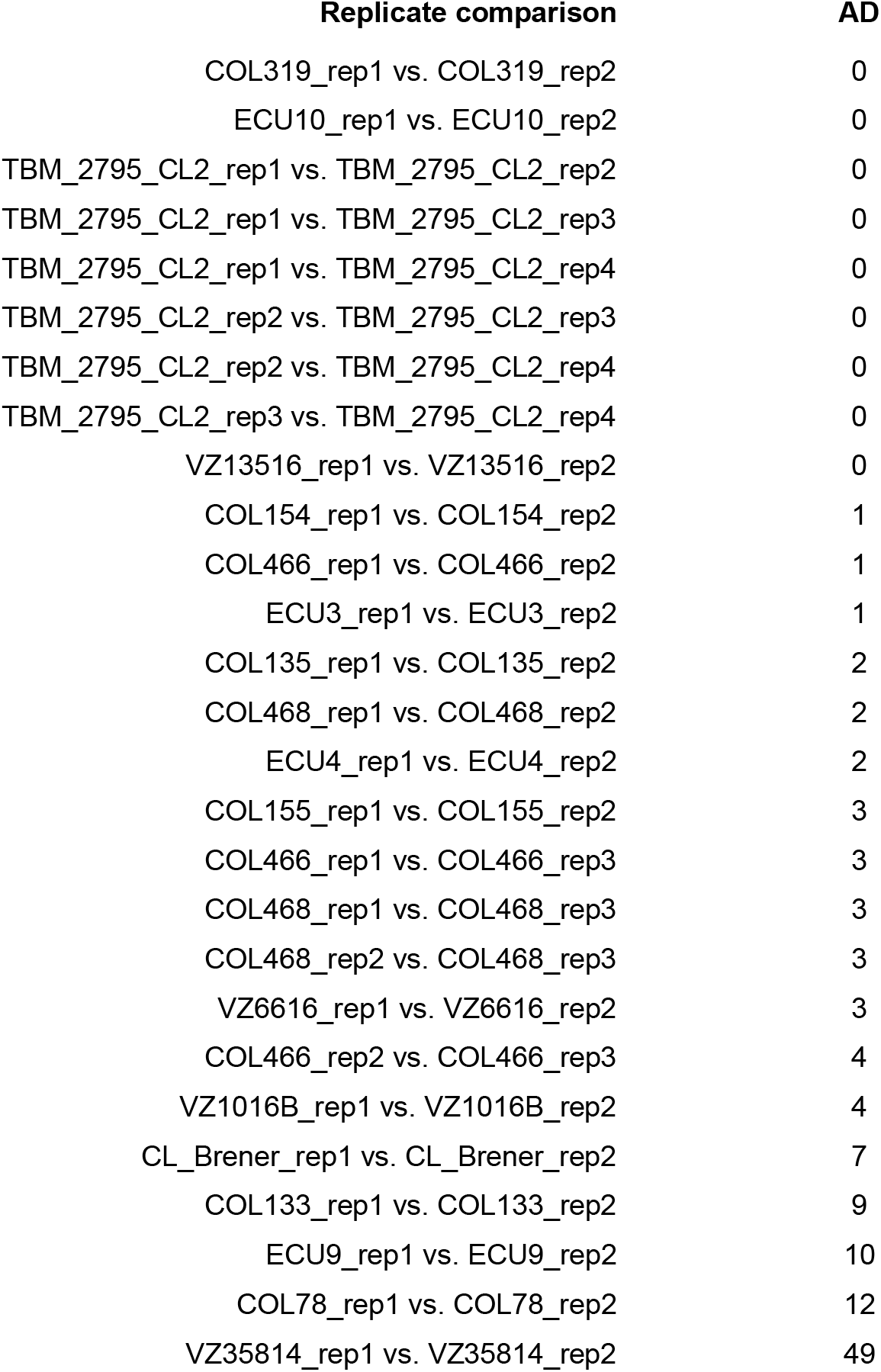
Allelic differences between GLST replicates. Eighteen samples were processed in 2 – 4 replicates after DNA extraction. A single SNP locus can differ by 0, 1 or 2 between two replicates (i.e., replicates can match at both, one, or neither allele). The AD measurement represents the total number of pairwise differences across all loci for which genotypes are called in all individuals (n = 561). The discrepancy between VZ35814 replicates likely represents barcode contamination with VZ16816 (see close similarity in Fig. 3).

**Table 2.** Basic diversity statistics for *T. cruzi* I samples from Colombia (COL), Venezuela (VZ) and Ecuador (ECU). Abbreviations: n (sample size); PS (polymorphic sites); HWE (Hardy-Weinberg equilibrium); F_IS_ (inbreeding coefficient), r^2^ (linkage coefficient), π (nucleotide diversity), Q (quartile); M (median); F_ST_ (between-group fixation index).

| Group (n) | PS | PS in HWE | $F_{IS}$ (Q1, M, Q3) | $r^2$ (Q1, M, Q3) | $\pi$ | $F_{ST}$ to COL | $F_{ST}$ to VZ | $F_{ST}$ to ECU |
| --- | --- | --- | --- | --- | --- | --- | --- | --- |
| COL (11) | 175 | 169 | -0.19, 0.13, 0.24 | 0.03, 0.07, 0.19 | 43.2 | 0.000 | 0.136 | 0.595 |
| VZ (7) | 147 | 143 | -0.35, -0.19, 0.11 | 0.02, 0.09, 0.27 | 29.0 | 0.136 | 0.000 | 0.632 |
| ECU (9) | 148 | 142 | -0.20, -0.09, 0.18 | 0.04, 0.17, 0.36 | 22.8 | 0.595 | 0.632 | 0.000 |

Genetic distances increased with spatial distances among samples (Mantel’s r = 0.89, p = 0.001), but the correlation coefficient was largely driven by high F_ST_ between sample sets from Colombia/Venezuela and Ecuador (Tbl. 2 and Fig. 5a): Mantel’s r decreased to 0.30 (p = 0.001) after restricting analysis to sample pairs separated by < 250 km (Fig. 5b). Within-country IBD appeared to grow stronger for samples separated by < 150 km (Mantel’s r = 0.48, p = 0.002) given a lack of correlation observed at higher distance classes within the Colombian dataset (Fig. 5b).

**Figure 5.**
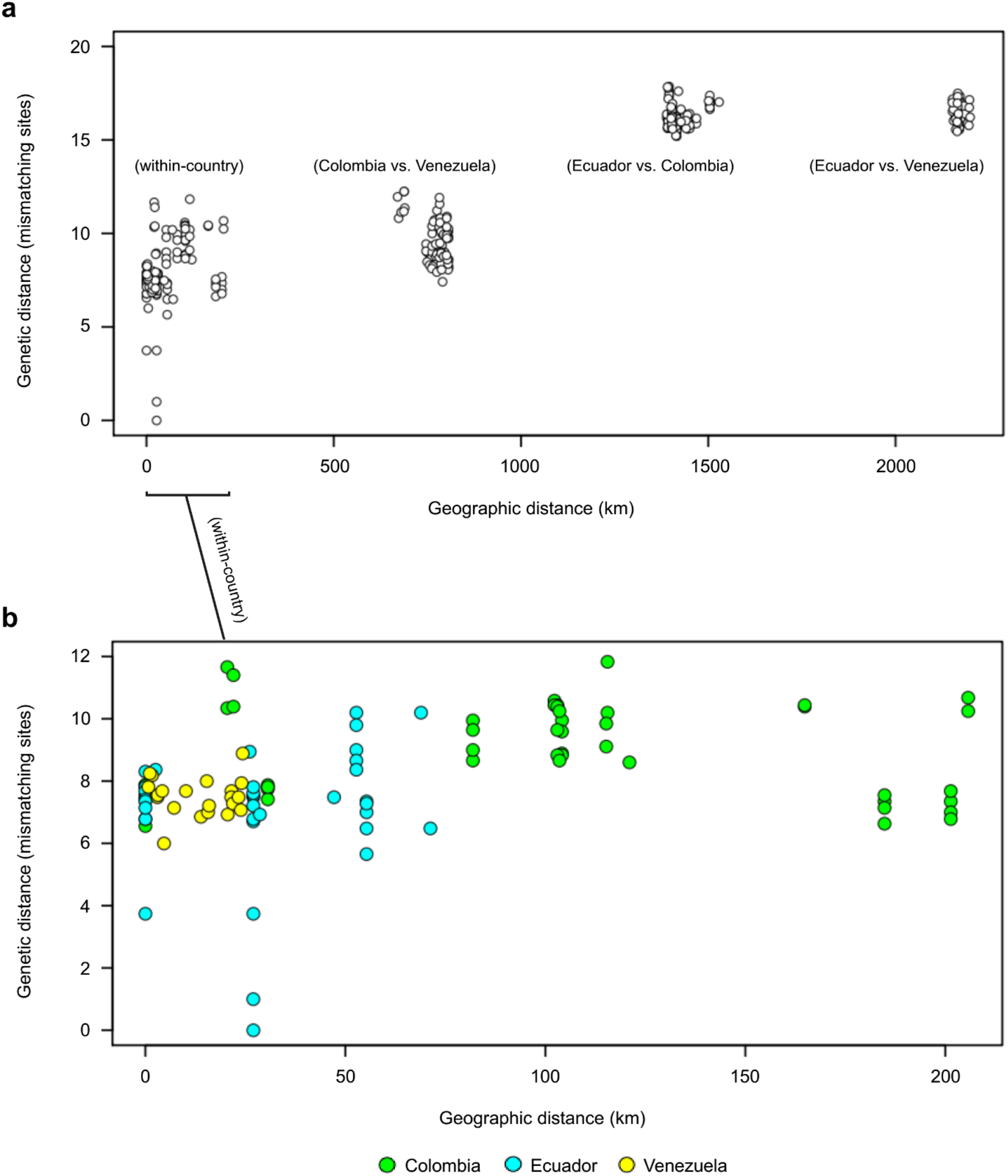
Isolation-by-distance among *T. cruzi* I samples. **a** Each circle represents geographic and genetic distances between two TcI samples. Global isolation-by-distance (IBD) is significant (Mantel’s r = 0.89, p = 0.001) but driven by divergence between Ecuadorian samples and the rest of dataset (see two clusters at top right). **b** Nevertheless, IBD remains significant for within-country comparisons at < 250 km (Mantel’s r = 0.30, p = 0.009) and < 150 km (Mantel’s r = 0.48, p = 0.002). Green, cyan and yellow fill colors represent comparisons within Colombia, Ecuador and Venezuela, respectively. Each of the above Mantel tests remains significant when sample pairs with genetic distances < 2 are removed (see arrows). Only variant sites with ≤ 10% missing genotypes (n = 285) are used in analysis. Only the first replicate is used for samples represented by multiple replicates.

Finally, GLST also clearly separated sub-lineages TcI, TcIII, TcIV, and TcVI in network (Fig. 3) and neighbor-joining tree construction (Fig. 6). AD between reference clones of different sub-lineages ranged from 153 (Arma18 cl1 (TcIV) vs. Para7 cl.3 (TcV)) to 472 (Chile c22 (TcI) vs. Saimiri3 cl. 8 (TcIV)).

**Figure 6.**
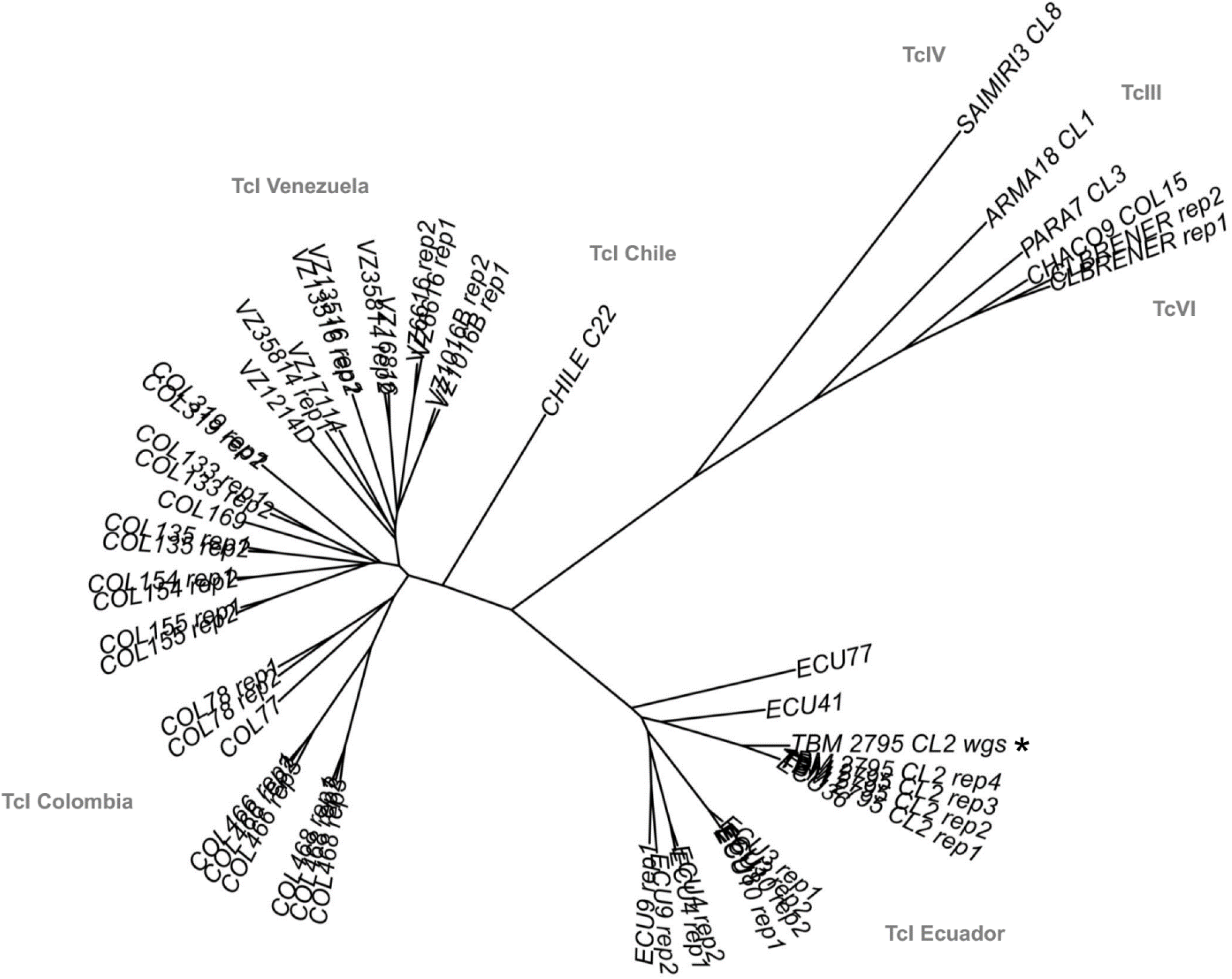
Neighbor-joining relationships among *T. cruzi* I samples and reference clones of other sub-lineages. Genetic distances are based on 556 biallelic SNP sites for which genotypes are called in all individuals. Results indicate high repeatability among most technical replicates (see ‘rep1 – 4’ suffices) and clearly separate TcI, TcIII, TcIV and TcVI. The tree also contains TBM_2795_CL2_wgs (see asterisk). This control sample was genotyped at the same 556 GLST loci using whole-genome sequencing (Illumina HiSeq) data from Schwabl et al. 2019^1^.

## Discussion

### Principle results

The GLST primer panel design and amplicon sequencing workflow outlined in this study aimed to profile *T. cruzi* genotypes at high resolution directly from infected triatomine intestinal content by simultaneous amplification of 203 genetic target regions that display sequence polymorphism in publicly available WGS reads. Mapped GLST amplicon sequences generated from *T. cruzi* reference clones and from metagenomic intestinal DNA extracts containing a minimum of 3.69 pg/μl *T. cruzi* DNA achieved high target specificity (< 1% off-target mapping) and yield (391 of 403 target SNP sites mapped). Mapping depth variation across target loci was highly repeatable between sequencing runs. 387 SNP sites were identified among *T. cruzi* DTU I samples and 393 SNP sites were identified in non-TcI reference clones. These markers showed low linkage and clearly separated *T. cruzi* individuals within and across DTUs, for the most part also individuals collected at the same or closely separated localities in Colombia, Venezuela, and Ecuador. An increase in pairwise genetic differentiation was observed with increasing geographic distance in analyses within and beyond 150 km.

### Cost-effective spatio-genetic analysis

GLST achieved an important resolution benchmark in recovering isolation-by-distance (IBD)^64^ at less than 150 km. These correlations indicate the potential of GLST in spatially explicit epidemiological studies which, for example, aim to identify environmental variables or landscape features that modify IBD^27^. High spatial sampling effort is typically required by such studies and often limits budget for genotyping tools. GLST appears promising in this context as library preparation costs < 4.00 USD per sample (see cost summary in Supplementary Tbl. 3) and can be completed comfortably in two days. The first-round PCR reaction requires very low primer concentrations (0.125 μM) such that a single GLST panel purchase (0.01 μmol production scale) enables > 100,000 reactions and can be shared by several research groups. Sequencing represents a substantial cost but is highly efficient due to short fragment sizes and few off-target reads. High library complexity also promotes the use of GLST in the role of PhiX, i.e., as a spike-in to enhance read quality in a different sequencing run. Our study easily decontaminated reads from a spiked amplicon pool sharing barcodes with GLST (run 1). Alternatively, i.e, when GLST is sequenced alone (run 2), one Illumina MiSeq run is expected to generate > 70x median genotype depth for 100 samples using Reagent Micro Kit v2 (ca. 1,000 – 1,500 USD, depending on provider; Supplementary Tbl. 3).

### GLST in relation to multi-locus microsatellite typing

We consider multi-locus microsatellite typing (MLMT) as the primary alternative for high-resolution *T. cruzi* genotyping directly from metagenomic DNA. MLMT has revolutionized theory on *T. cruzi* ecology and microevolution, for example, on the role of disparate transmission cycles^65,66^, ecological host-fitting^67^ and ‘cryptic sexuality’^68^ in shaping population genetic structure in TcI. In some cases^69,70^ (but others not^66,67,71^), the hypervariable, multiallelic nature of microsatellites allows every sample in a dataset to be distinguished with a different multi-locus genotype (MLG). This depends on panel size and spatial scale but also on local reproductive modes – e.g., sampling from clonal sylvatic vs. non-clonal domestic transmission cycles has correlated with the presence or absence of repeated MLGs^66^. In this study, we found two identical GLST genotypes shared among five samples from southern Ecuador. All other samples appeared unique, including those from Venezuela, where triatomine collection occurred at seven domestic localities within the city of Caracas. The small subset of repeated genotypes found in this study may reflect patchy, transmission cycle-dependent clonal/sexual population structure in southern Ecuador (see Schwabl et al. 2019^1^ and Ocaña-Mayorga et al. 2010^66^) but may also represent a weakness in GLST compared to MLMT in tracking individual parasite strains. The use of large MLMT panels, however, is significantly more resource-intensive because each microsatellite marker requires a separate PCR reaction and capillary electrophoresis cannot be highly multiplexed. MLMT data are poorly archivable across studies and may also be less suitable for inter-lineage phylogenetic analyses due to unclear mutational models and artefactual similarity from saturation effects^72^. Although our GLST panel was designed for TcI, its focus on syntenous sequence regions enabled efficient co-amplification of non-TcI DNA. GLST clearly separated TcI samples from all non-TcI reference clones, with highest divergence observed in Saimiri3 cl. 8. Interestingly, large MLMT panels have shown comparatively little differentiation between this sample and TcI, also more generally suggesting that TcIV and TcI represent monophyletic sister clades^72^.

### Adjustment and transferability

Considering the great variety of sample types to which studies have successfully applied PCR^73–77^, we expect that GLST can be applied to metagenomic DNA from many host/vector tissue types, not only from triatomine intestine as shown here. Further tests are required to determine whether low *T. cruzi* DNA concentrations in chronic infections or sparsely infected organs (e.g., liver and heart^78^) are also amenable to GLST. We focused analysis on *T. cruzi* DNA concentrations of at least one picogram per microliter metagenomic DNA (this equates to ca. 30 parasites per microliter in the case of TcI^79^) without heavily investigating options to enhance sensitivity or sensitivity measurement, for example, by additional removal of PCR inhibitors, improved primer purification (e.g., HPLC vs. salt-free), post-PCR probe-hybridization^80^ or barcoding/sequencing of samples with unclear first-round PCR amplicon bands. Even relatively aggressive processing methods may be tolerable given that DNA fragmentation is unlikely to compromise the 120 – 160 nt size range targeted by GLST. Increasing sensitivity by increasing PCR amplification cycles, however, is less advised. PCR error appeared relevant with as little as 30x (+ 7x barcoding) amplification in this study as we observed noise among replicates despite high read-depth and SNP-call overlap between sequencing runs. Rates or error were, however, well within margins expected for methods involving PCR^81^. We also note that the exceptional discrepancy between VZ35814 replicates unlikely represents systematic error but barcode contamination with VZ16816. Such error is perhaps less likely if primers are kept in separate vials instead of in the plate format which we have used here.

Wet lab aside, the main objective of this study was to provide a transparent bioinformatic workflow for highly multiplexable primer panel design using freely available softwares and publicly archived WGS reads (e.g., see www.ebi.ac.uk/ena or www.ncbi.nlm.nih.gov/sra). Importantly, we show that knowledge of polymorphic genetic regions in parasite genomes from one small study area (Loja Province, Ecuador) can suffice to guide variant discovery at distant, unassociated sampling sites. Our demonstration using *T. cruzi* should be easily transferable to any other pathogenic species with a published reference genome. Target selection can also be tailored to a variety of objectives. For example, while landscape genetic studies on dispersal often focus on neutral or non-coding sequence variation^82^, experimental (e.g., drug testing) studies may seek to detect single-nucleotide changes in coding regions, perhaps in genes belonging to specific ontology groups or associated with results of high-throughput proteomic screens^83^. The candidate SNP pool can easily be filtered for such criteria during GLST panel design, e.g., using SnpEff^84^ or BEDTools^85^ and data mining strategies at EuPathDB^86^. Candidate SNP filtering by minor allele frequency (MAF) may also be useful when the target population is closely related to that of the WGS dataset guiding panel design. Placing a minimum threshold on MAF (using VCFtools^87^, etc.), for example, may improve analyses of population structure and genealogy whereas a focus on low-frequency variants may help in tracking individuals or recent gene flow at the landscape scale^88^. It may also be possible to refine panel design towards markers that meet model assumptions in later analysis. Hardy Weinberg Equilibrium (HWE), for example, is a common requirement in demographic modelling^89–91^, Bayesian clustering^92^, admixture/migration^93,94^ and hybridization tests^95^. Deviation from HWE may occur more frequently in specific genetic regions (e.g., near centromeres^96^), and SNPs in these could be excluded from the target pool. Numerous other filtering options – e.g., based on allele count (to enhance resolution per SNP), distance to insertion-deletions (to improve target alignment), or percent missing information (to avoid poorly mapping regions) – are easily implemented with common analysis tools^97^.

GLST is also highly scalable because increasing panel size does not lead to more laboratory effort or processing time. Sequencing depth requirements and thermodynamic compatibilities among primers are more relevant in limiting panel size. However, it is also possible to divide large GLST panels into two or more PCR multiplexes based on ΔG-based partitioning in MultiPLX^98^. Unintended primer affinities (i.e., polymer formations) can also be removed by gel excision, e.g., as we have done using the PureLink Quick Gel Extraction Kit.

### Prospects

This study sought to provide a framework for various epidemiological research but was restricted in its own ability to make important inferences on *T. cruzi* ecology because only few samples (remainders from different projects) were analyzed. Samples were also aggregated either to domestic or to sylvatic ecotopes (see Supplementary Tbl. 1). More extensive, purposeful sampling could have, for example, helped us explore whether COL468’s position deep within the Cordillera Oriental contributes to its strong divergence to samples such as COL135 or COL319, these perhaps more closely related due to lower ‘cost-distances’^99^ along the basin range. Fuelling landscape genetic simulators such as CDMetaPOP^91^ with high GLST sample sizes is an especially exciting direction for future research. It would also be interesting, for example, to extend this study’s sampling to cover gradients along the perimeter of Caracas and adjacent El Ávila National Park (see Fig. 4b). Sylvatic *P. geniculatus* vector populations appear to be rapidly adapting to habitats within Caracas^39,100^ but parallel changes in the distribution of *T. cruzi* genetic diversity have yet to be tracked. The low cost of GLST also makes it more feasible for studies to simultaneously assess genetic polymorphism in each vector individual from which parasite markers were amplified. Such coupled genotyping would enhance resolution of parasite-vector genetic co-structure and thus, for example, help quantify rates of parasite transmission from domiciliating vectors or determine whether parasite gene flow proxies for (or improves understanding of) dispersal patterns in more slowly evolving vectors or hosts. It would also be interesting to test in how far deep-sequenced GLST libraries could help in detecting (and reconstructing distinct MLGs from) multiclonal *T. cruzi* infections without the use of cloning tools^101^, e.g., using bioinformatic strategies developed for malaria research^102–105^. Multiclonality has important implications for public health^106,107^ but its potential prevalence in *T. cruzi* vectors and hosts^108,101,109^ is difficult to describe from cultured cells^108,110^. Countless other applications are conceivable for GLST. Some research fields, however, will surely be less amenable to the PCR-based approach. Relative amplicon concentrations, for example, appeared to be too stochastic in this study to allow inference of copy number variation or other structural rearrangements based on read-mapping depths. Unintended primer alignment is also likely to occur if PCR targets are located within highly repetitive sequences such as those encoding surface protein families in sub-telomeric regions of the *T. cruzi* genome^46^.

We look forward to seeing GLST approaches in a wide variety of research for which such limitations do not apply. Regarding population and landscape genetic studies, prudent spatial and genetic sampling design is often key to meaningful inference and we hope that the low cost and high flexibility of our pipeline helps researchers achieve all criteria required.

## Supplementary figures and tables

**Supplementary Figure 1.**
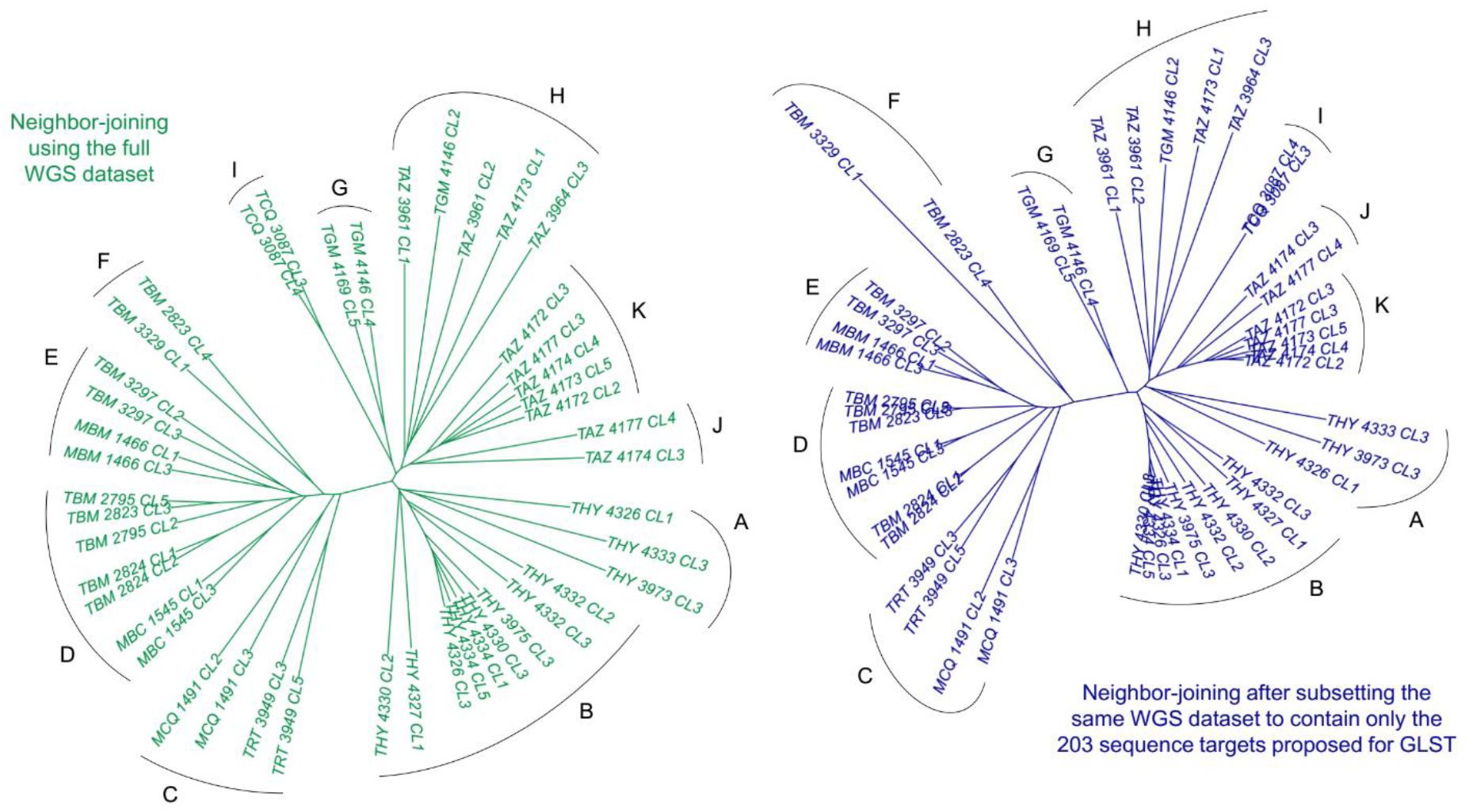
Phylogenetic resolution at GLST loci *in silico*. The green tree shows neighbor-joining (NJ) relationships calculated from 106,007 SNP sites identified from whole-genome sequencing (WGS) of 45 TcI clones in southern Ecuador^1^. Sites missing genotypes in ≥ 10% individuals are excluded. Less than 45 km separate the most distant sampling sites within the study region. Several pairs of clones also represent the same host/vector individual (see first seven characters of IDs). NJ was repeated after abridging the WGS dataset to contain only SNPs within the 203 sequence targets proposed by GLST (also excluding sites missing ≥ 10% genotypes). This resultant tree (blue, at right) uses 391 SNP sites and recreates clusters A-K observed in WGS.

**Supplementary Figure 2.**
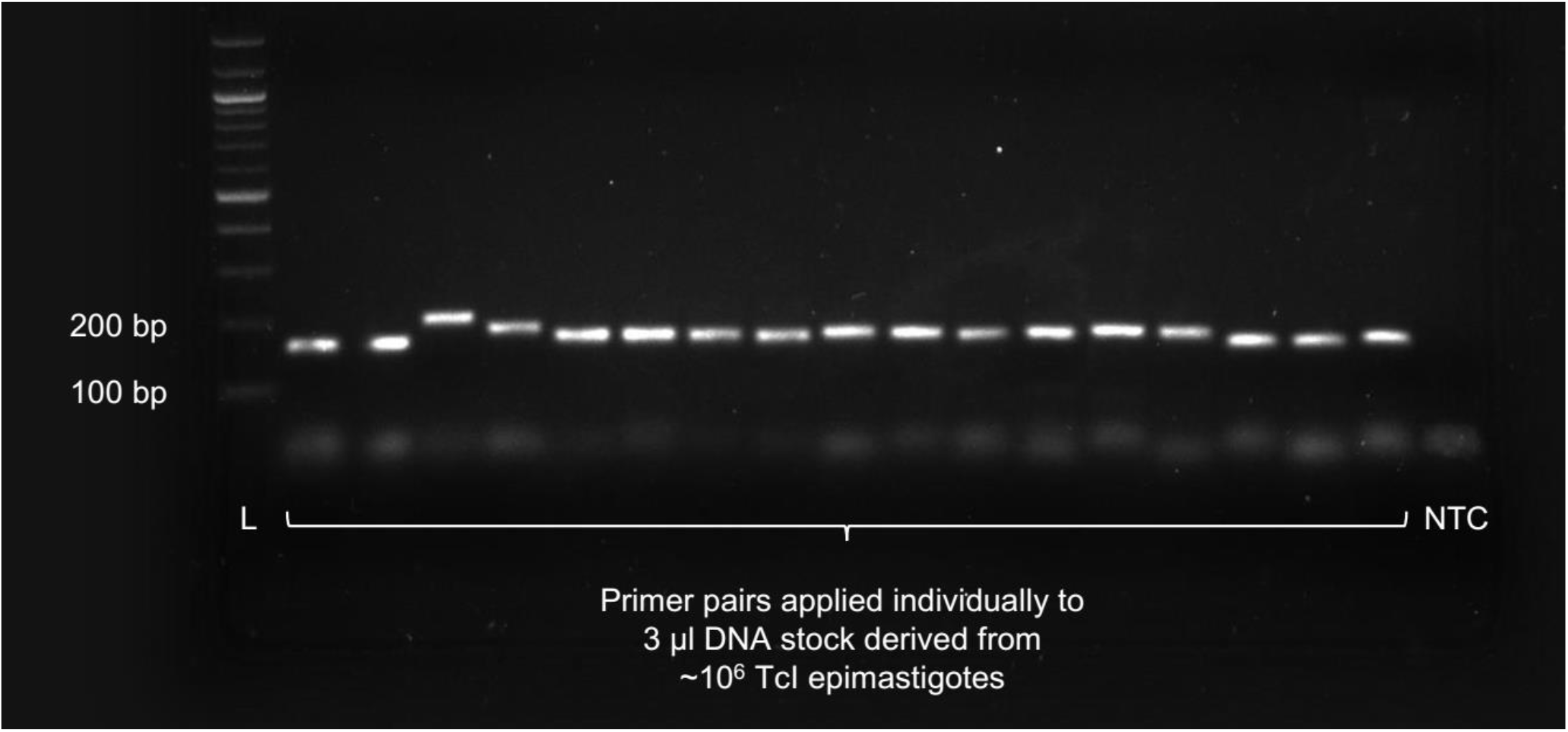
Individual primer pair validation. Primer pairs were first applied individually to pure TcI epimastigote DNA to confirm product amplification within the expected size range (164 – 204 bp). The figure shows the electrophoresed products of 17 different primer pairs in 0.8% agarose gel as well as DNA ladder (L) and no-template control (NTC). All other primer pairs achieved similar results using an initial incubation step at 98 °C (2 min); 30 amplification cycles at 98 °C (10 s), 60 °C (30 s), and 72 °C (45 s); and a final extension step at 72 °C (2 min).

**Supplementary Figure 3.**
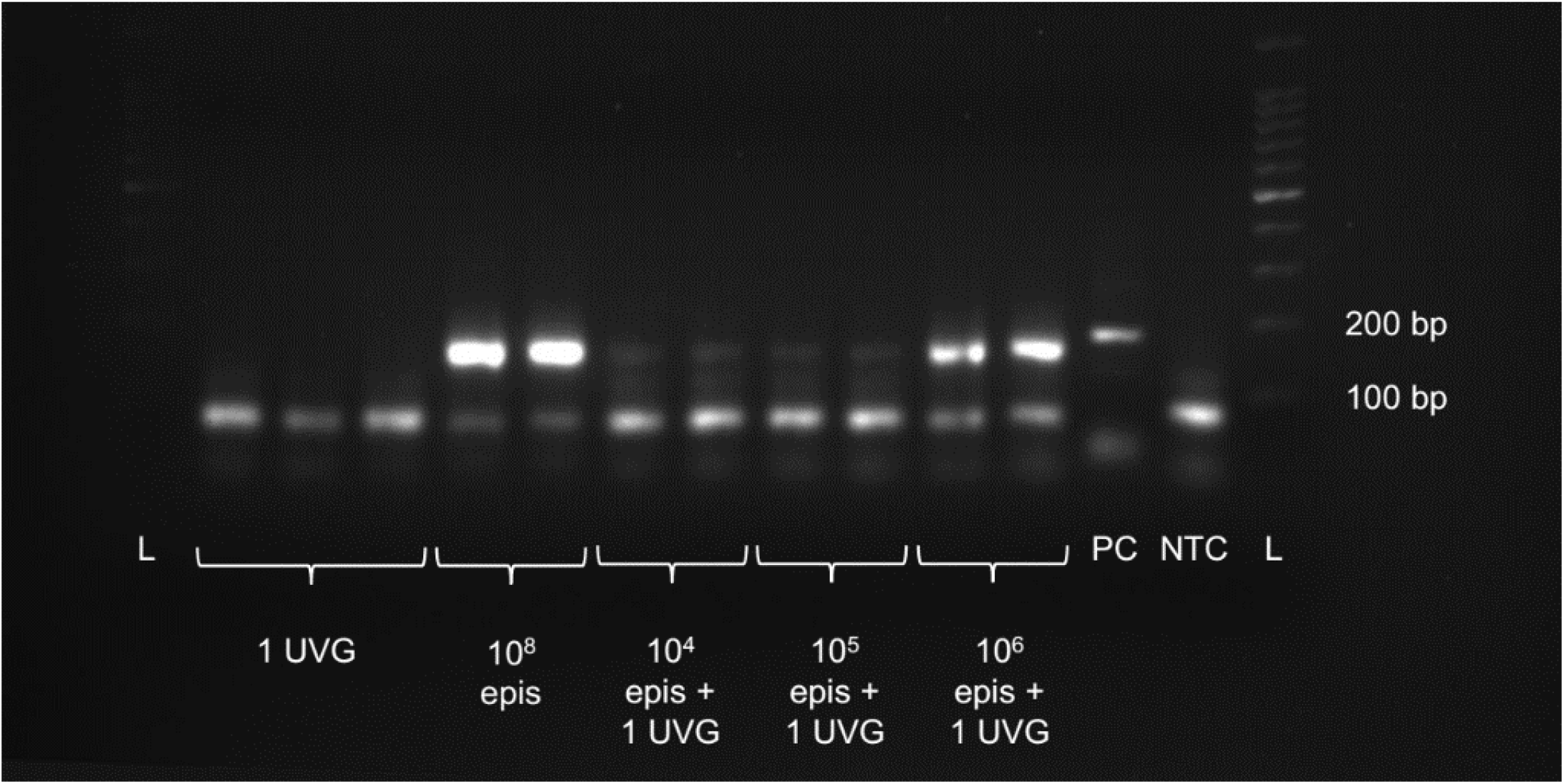
Preliminary GLST (multiplex) trials on *T. cruzi* I mock infections. We created mock infections by mixing 10^4^, 10^5^ and 10^6^ RNAlater-preserved TcI-Sylvio epimastigote (epi) cells with uninfected *Rhodnius prolixus* vector gut (UVG). DNA extracted from these mock infections was subjected to the multiplexed, 203-target GLST reaction (using the same cycling conditions as for single-target reactions – see Methods or Supplementary Fig. 2 legend) and products were electrophoresed in 0.8% agarose gel. Fainter banding of GLST products from lower concentration mock infections encouraged follow-up on sensitivity thresholds using additional dilution curves and qPCR. Next to DNA ladder (L) and no-template control (NTC), the gel also contains TcZ primer product from pure TcI epimastigote DNA. TcZ primers provide a highly sensitive positive control (PC) as they target 195 bp satellite DNA repeats that make up ca. 5% of the *T. cruzi* genome.

**Supplementary Figure 4.**
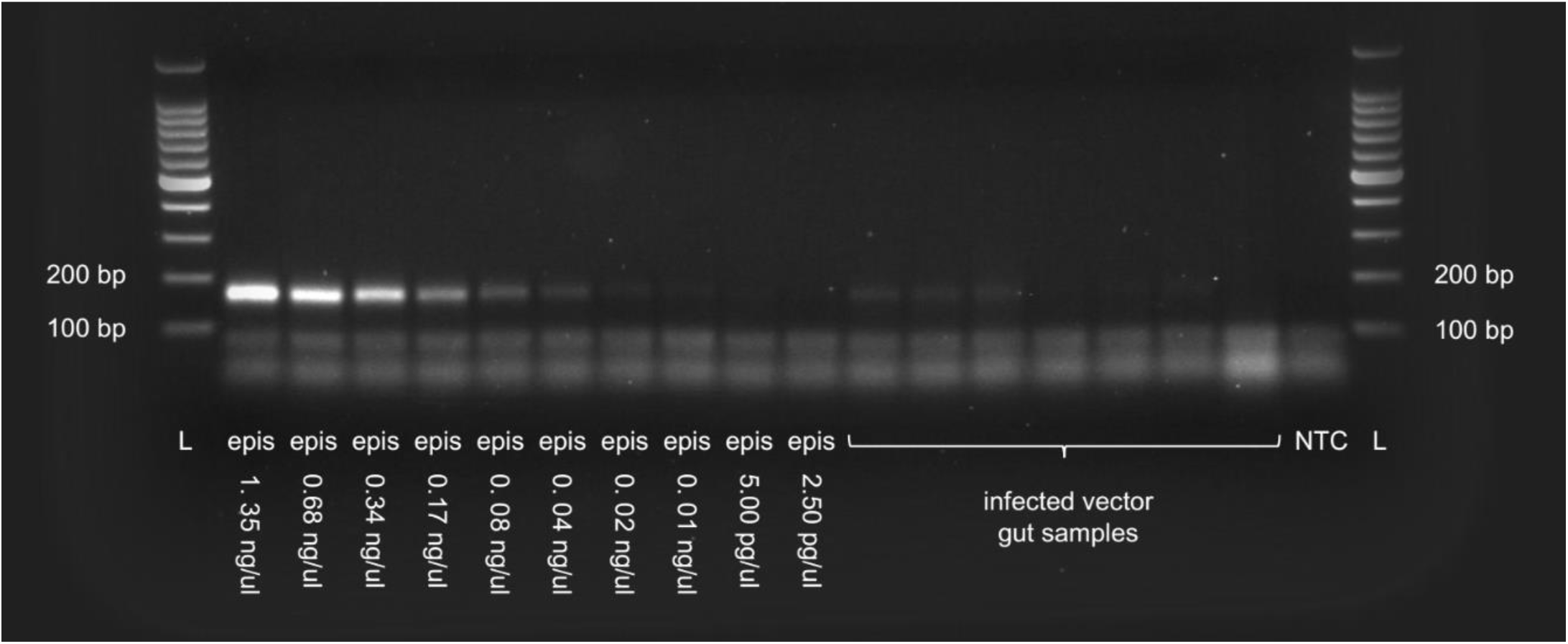
*T. cruzi* I DNA dilutions and GLST product visibility in 0.8% agarose gel. The left side shows electrophoresed GLST amplicons generated from 3 μl pure TcI epimastigote (epi) DNA with concentrations between 1.35 ng/μl and 2.50 pg/μl (see cycling conditions in Methods or Supplementary Fig. 2 legend). Lanes on the right contain amplicons from seven random metagenomic samples that tested positive for *T. cruzi* satellite DNA (not shown). DNA ladders (L) and no-template control (NTC) are indicated left and right. Poor amplicon visibility occurs at ≤ 60 pg epimastigote DNA input. Gut DNA amplicon visibility is also limited but whether this relates to low *T. cruzi* content or amplification interference is unclear without qPCR.

**Supplementary Figure 5.**
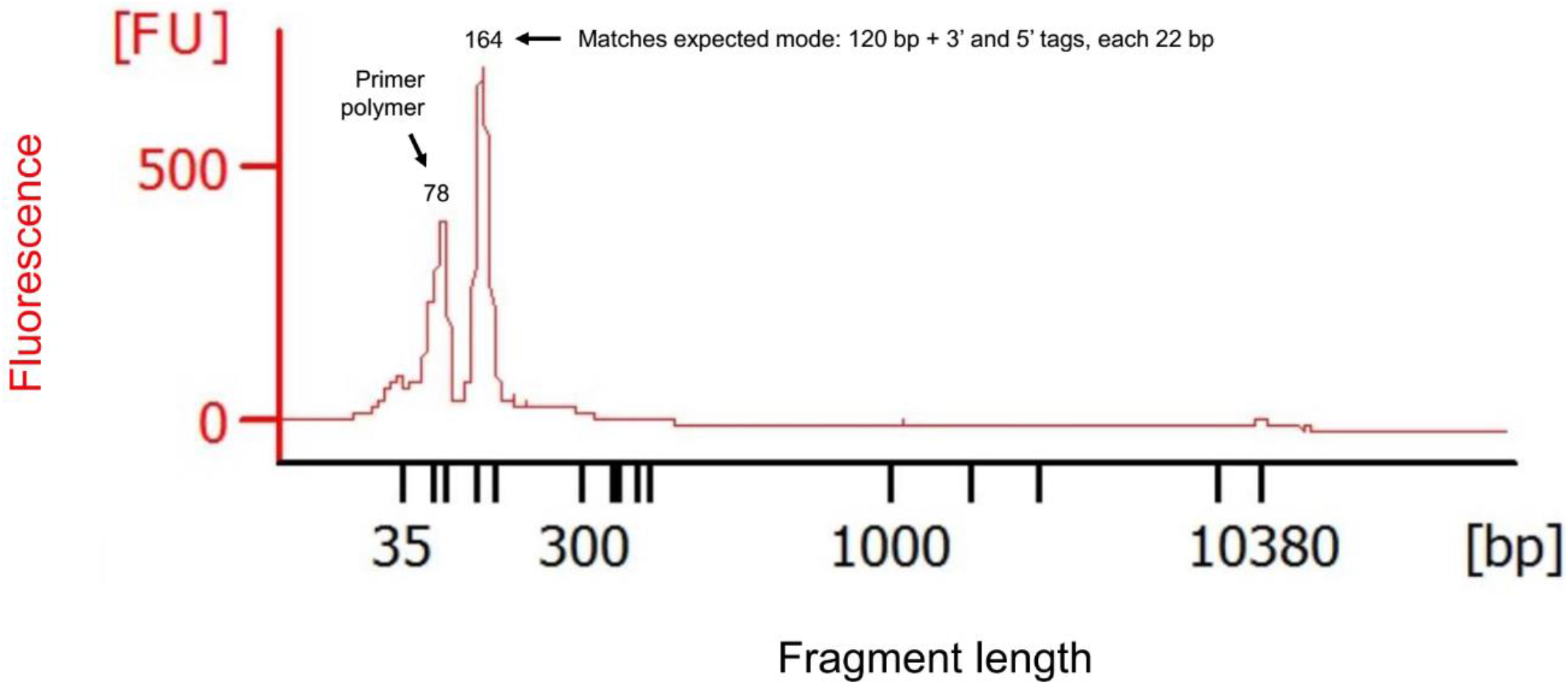
First-round (unbarcoded) PCR product size composition measurement using microfluidic electrophoresis. The figure plots fragment sizes (calculated based on migration times relative to those of standards) and fluorescence intensity (FU) of first-round PCR products (see cycling conditions in Methods or Supplementary Fig. 2 legend) measured with the Agilent Bioanalyzer 2100 System. The first peak represents primer polymerization that is removed in subsequent gel excision/re-solubilization steps. The second peak matches expectations for the multi-target GLST product (164 – 204 bp). Special thanks to Craig Lapsley at the Wellcome Centre for Molecular Parasitology in Glasgow for generating this data.

**Supplementary Figure 6.**
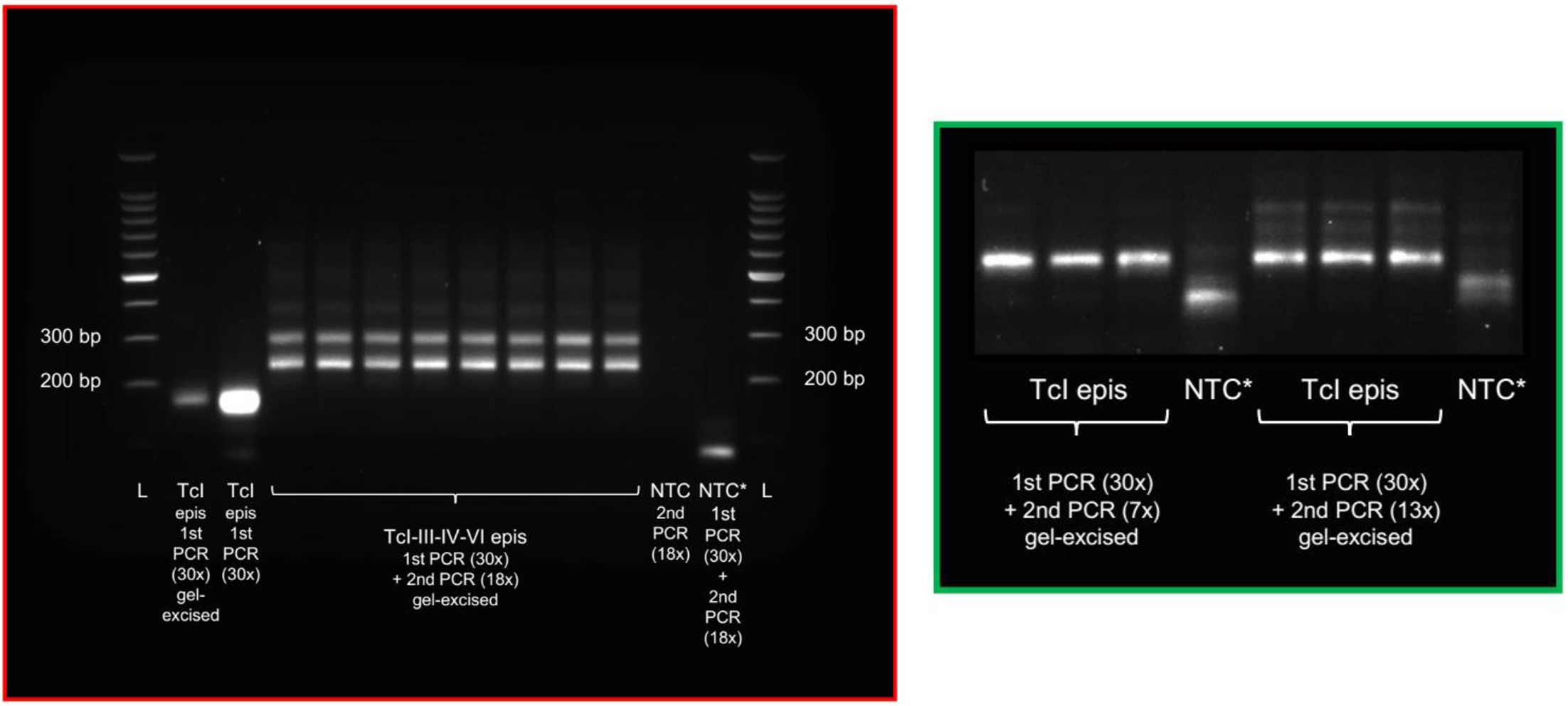
Large polymer formation from excessive amplicon barcoding. The second (barcoding) PCR reaction uses an initial incubation step at 98 °C (2 min); 7 amplification cycles at 98 °C (30 s), 60 °C (30 s), and 72 °C (1 min); and a final extension step at 72 °C (3 min). Seven amplification cycles were chosen because unwanted polymers formed at 13 and 18x. The center lanes in the 0.8% agarose gel at left (red border) show electrophoresed GLST products from reference clones after eighteen cycles of barcoding PCR. Large, non-target banding occurs at ≥ 300 bp. Unbarcoded products from TcI epimastigote (epi) DNA are also shown at left. No template controls from barcoding (NTC) and first-round + barcoding PCR (NTC*) occur next to the DNA ladder (L) on the right side of the gel. The smaller image (green border) to the right shows how unwanted banding becomes less pronounced at 13x and largely disappears at 7x. This 0.8% agarose gel also contains NTC* samples, i.e., negative controls carried through both first and second-round PCR.

**Supplementary Figure 7.**
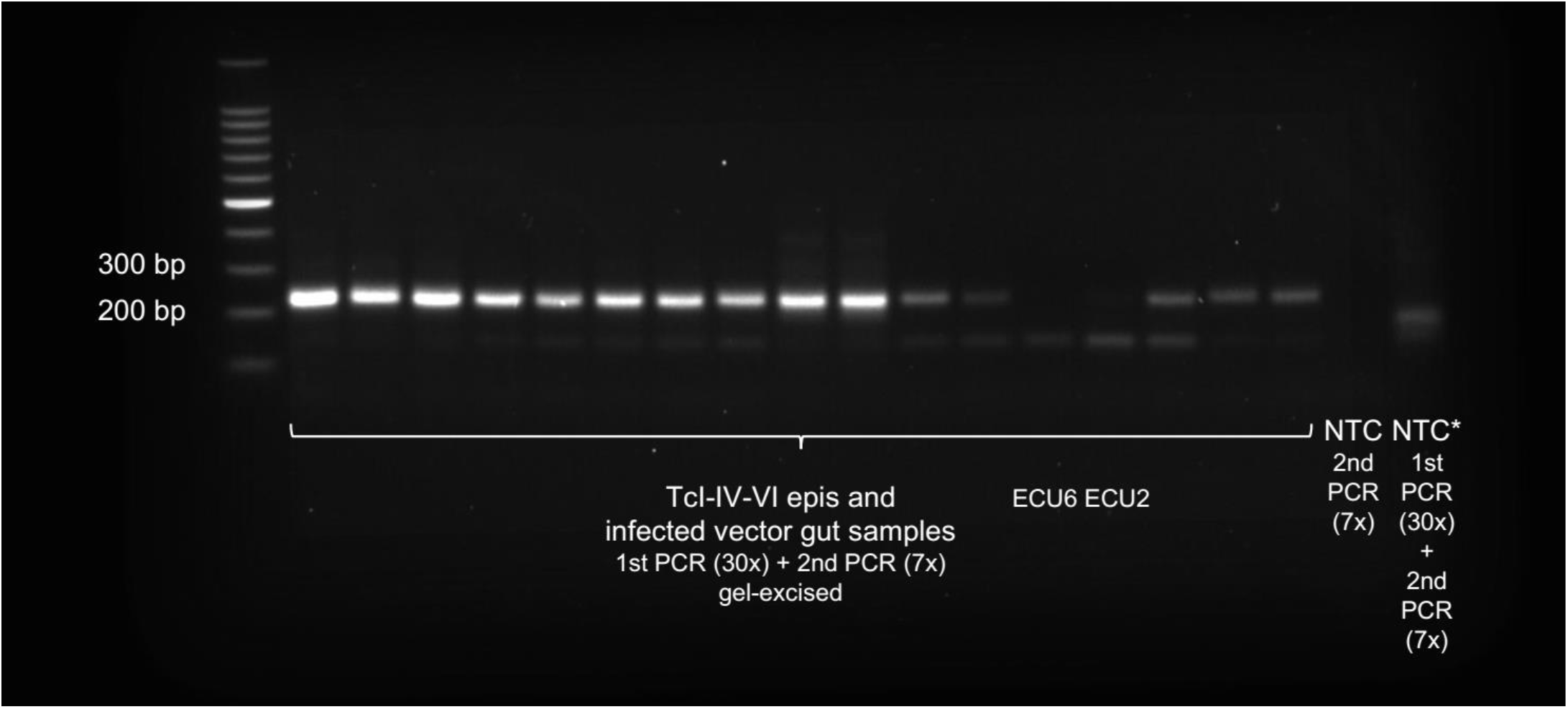
Barcoded GLST products ready for final pooling and purification. The 0.8% agarose gel shows a subset of fifteen GLST products from the second-round (barcoding) PCR reaction (see cycling conditions in Methods or Supplementary Fig. 6 legend) prior to equimolar pooling and final gel excision/re-solubilization steps. Products from ECU6 and ECU2 occur in this gel but were not included in the final pool. The gel also contains DNA ladder (L) and no-template controls from barcoding (NTC) and first-round + barcoding PCR (NTC*).

**Supplementary Figure 8.**
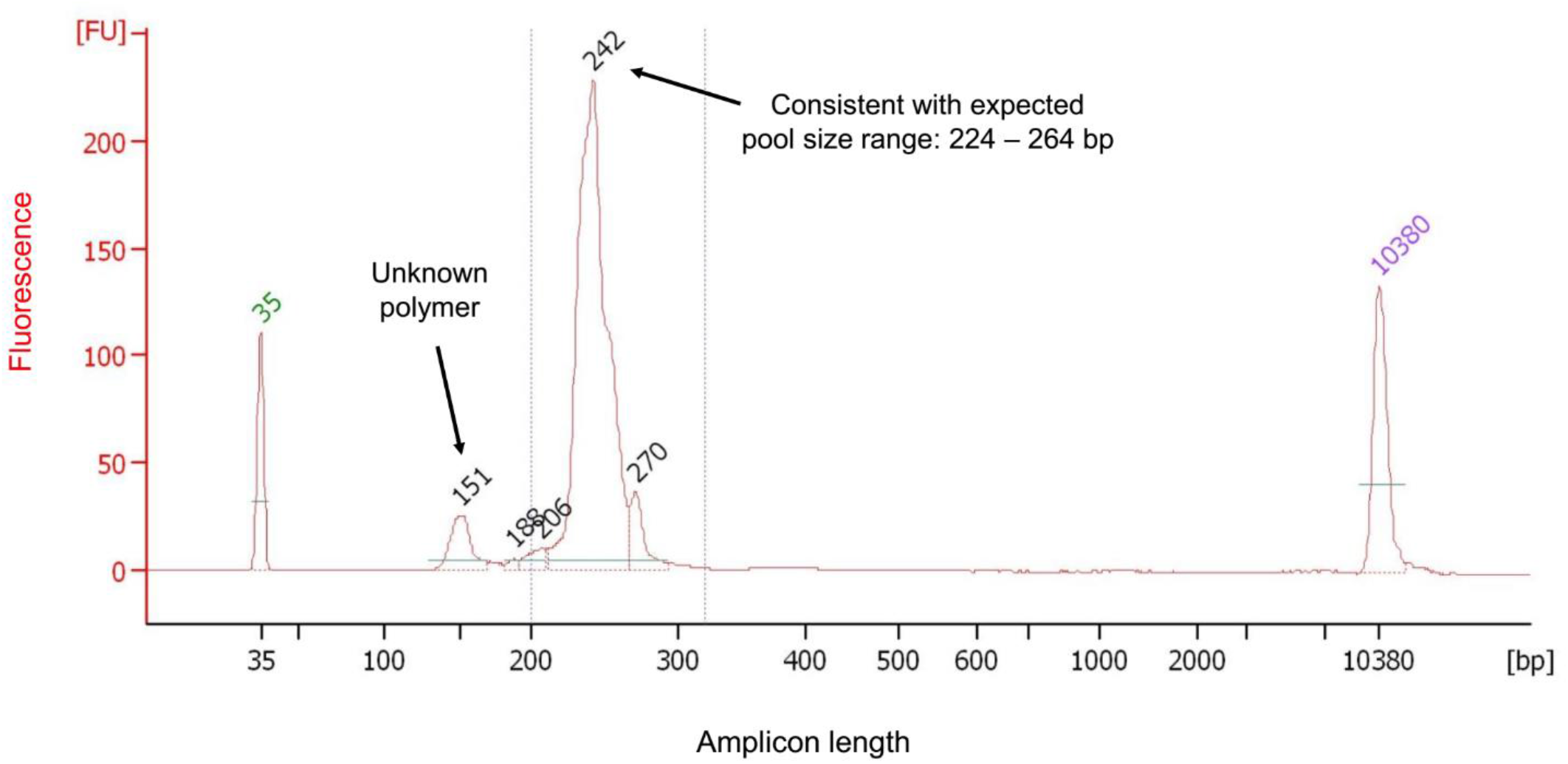
Final (barcoded) GLST pool size composition measurement using microfluidic electrophoresis. The figure plots fragment sizes (calculated based on migration times relative to those of standards) and fluorescence intensity (FU) of the final GLST pool measured with the Agilent Bioanalyzer 2100 System. The large peak matches expectations for the multi-target GLST product pool (224 – 264 bp). Left and right peaks labelled in green and purple represent standards of known size. A small non-target peak remaining near 151 bp encourages improvement of prior size selection steps. Special thanks to Julie Galbraith at Glasgow Polyomics for generating this data.

**Supplementary Figure 9.**
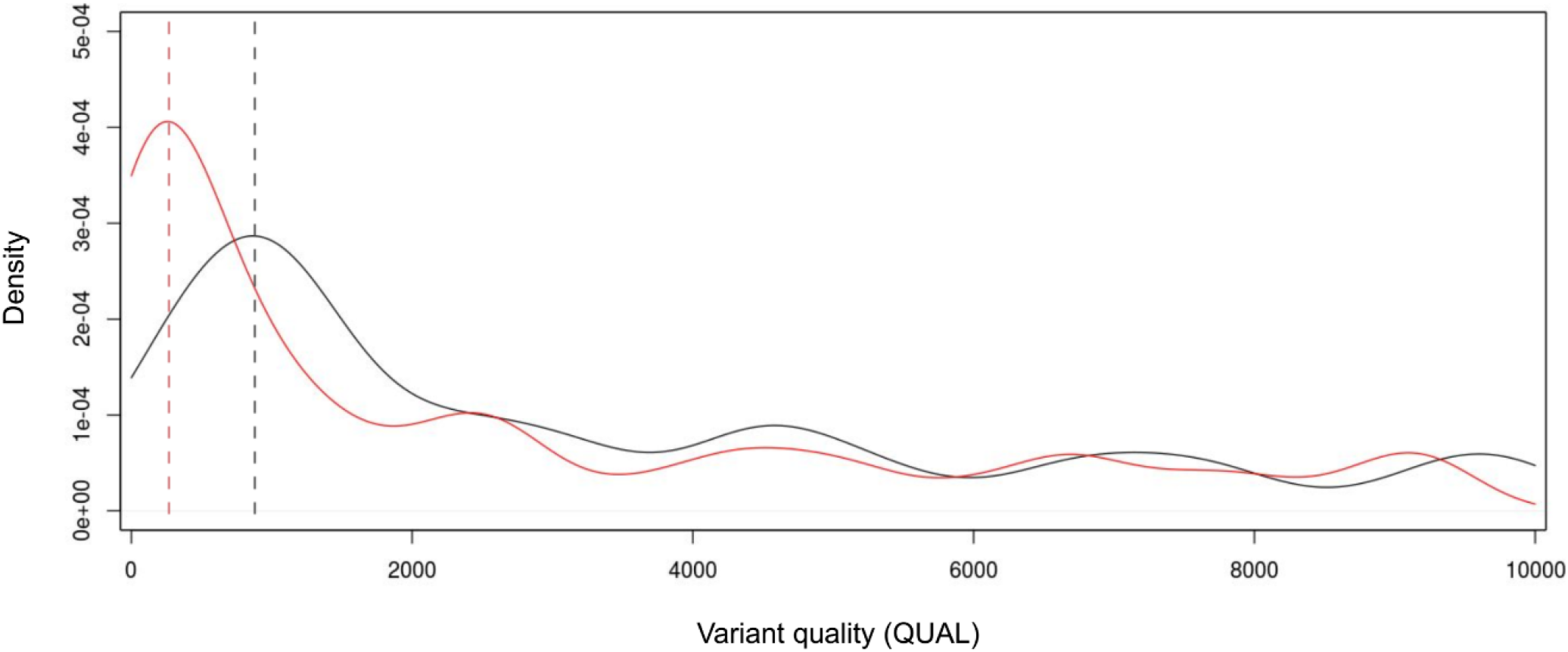
Quality scores at previously identified vs. unidentified variant sites. The GLST primer panel was designed based on single-nucleotide polymorphisms (SNPs) in Ecuadorian TcI clones. It was applied, however, to samples from distant geographic locations as well as to non-TcI clones. Additional, previously unidentified SNP sites (PU) were thus expected to be found but we needed to distinguish true PU from PCR and sequencing error. We reasoned that quality statistics (e.g., mapping quality, strand bias, minor allele frequency, etc. – see Methods) at previously identified SNP sites (PI) could help calibrate quality filters applied to the wider dataset. This strategy finds support in the above density plot of QUAL scores computed by Genome Analysis Toolkit^42^. The plot suggests that, prior to variant filtration, lower QUAL scores occur more often at PU (red) than at PI (black). We thus imposed the most stringent filtering criteria possible without losing PI.

**Supplementary Figure 10.**
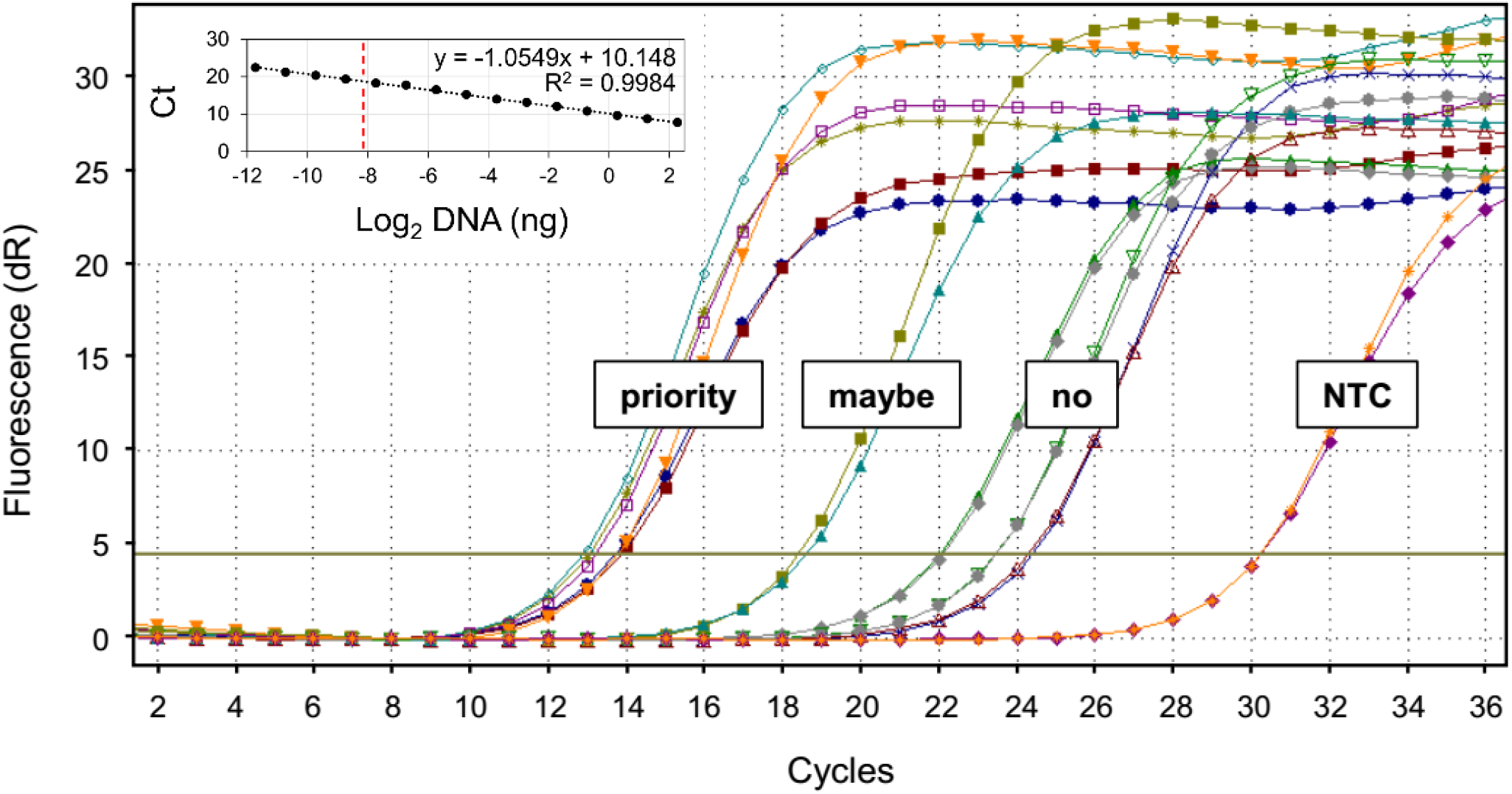
GLST sample selection and sensitivity estimation via qPCR. We used *T. cruzi* satellite DNA qPCR to identify vector gut samples with *T. cruzi* DNA quantities within ranges successfully visualized in GLST reactions using epimastigote DNA (Supplementary Fig. 4). The qPCR reaction used an initial incubation step at 95 °C (10 min) and 40 amplification cycles at 95 °C (15 s), 55 °C (15 s), and 72 °C (15 s). The plot shows baseline-corrected fluorescence (dR) for seven sample duplicates. Following the regression equation from the standard curve (see inset), the three samples with highest cycle thresholds (Ct values) in this example represent gut extracts with 0.05 to 0.14 ng/μl *T. cruzi* DNA. Such samples with *T. cruzi* DNA concentrations above 0.01 ng/μl were prioritized for GLST and none failed in library construction. ECU36, with a mean Ct value of 18.68 in the plot, was also successfully sequenced. A Ct value of 18.68 represents 3.69 pg/μl *T. cruzi* DNA. Not all samples with concentrations at single-digit picogram levels (per μl) were successful and we did not troubleshoot those with substantially lower concentrations based on qPCR.

**Supplementary Figure 11.**
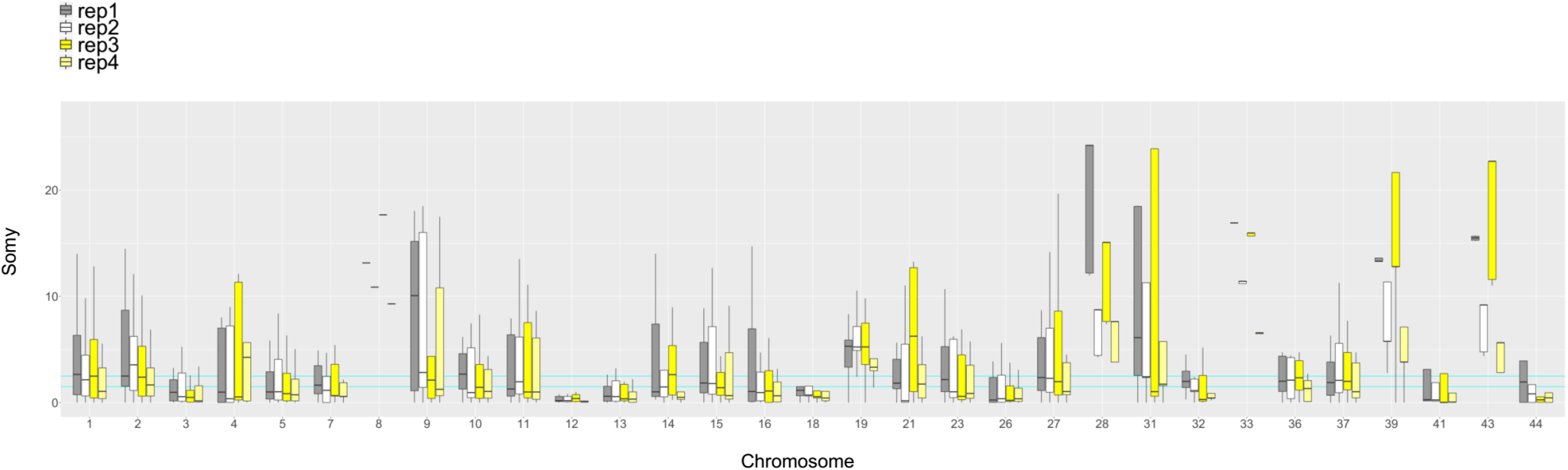
Target coverage in control replicates confirms expectations that the GLST panel applied in this study is unreliable for copy number estimation. We adapted methods from Schwabl et al. 2019^1^ to derive somy estimates for each base position within GLST amplicons. Briefly, we calculated median-read-depth of all target bases for each chromosome. We let the median of these chromosomal medians (the ‘inter-chromosomal median’) represent expectations for the disomic state, estimating copy number per base position by dividing each position’s read-depth by the inter-chromosomal median and multiplying by two. Boxplots show median and interquartile ranges of these site-wise somy estimates for each chromosome in TBM_2975_CL2 control replicates. TBM_2795_CL2 did not show chromosomal amplifications in whole-genome analysis^1^. Not unexpectedly for a PCR-based method, somy values estimated from GLST read-depths differ substantially among replicates and are unrealistically high/low on many chromosomes. Estimates on chromosomes with few GLST targets appear especially unreliable – e.g., see chromosomes 8, 28, 33, 39 and 43. These chromosomes contain ≤ 2 GLST targets each. The horizontal lines cyan lines mark y = 1.5 and y = 2.5.

**Supplementary Figure 12.**
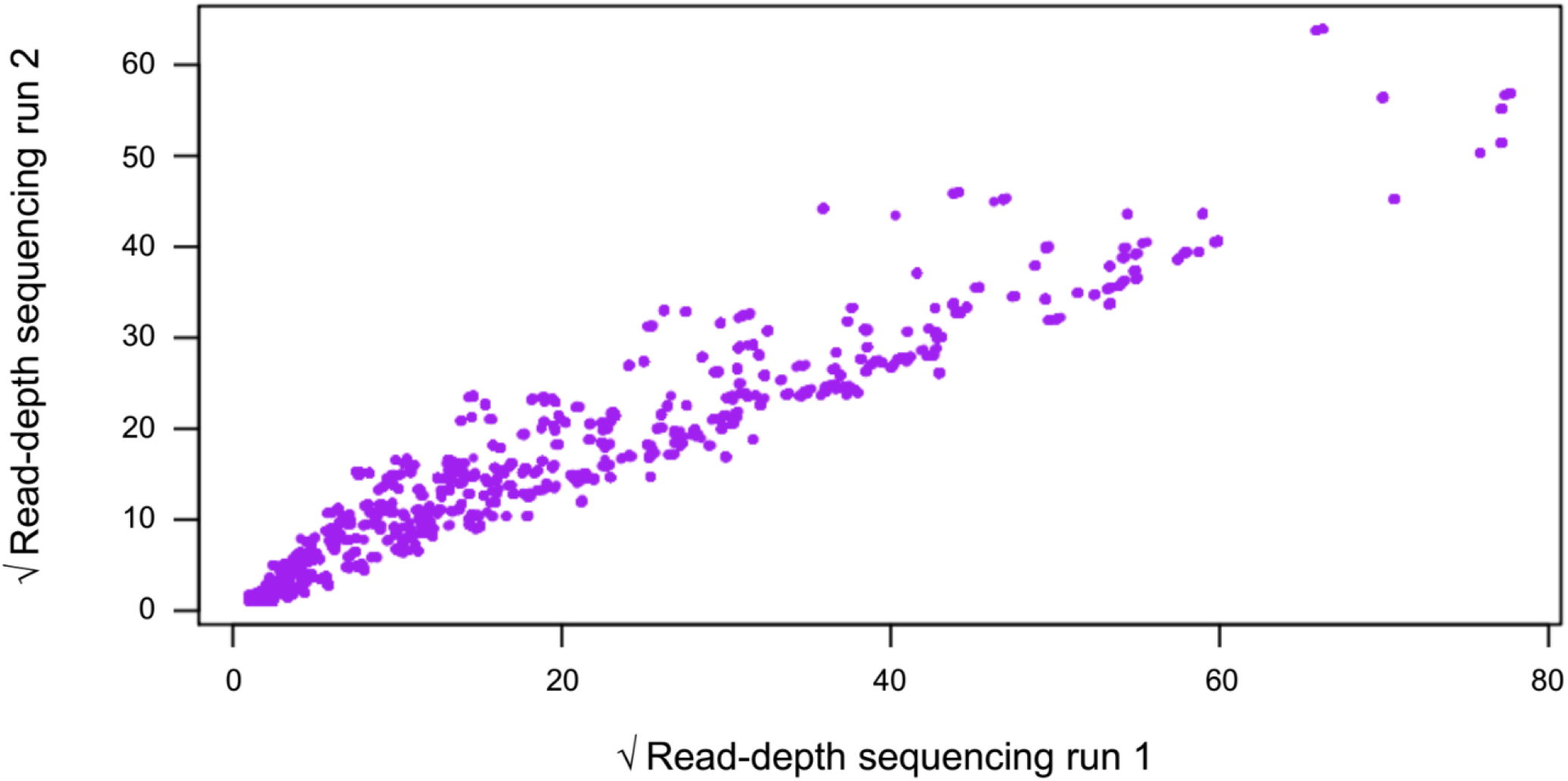
Similar read-depth distribution between separate sequencing runs. We sequenced the same GLST pool in two separate Illumina MiSeq runs. Run 1 involved GLST as a spike to a collaborator’s 16S amplicon library, whereby GLST reads were subsequently decontaminated from (barcode-sharing) 16S reads by alignment to the TcI-Sylvio reference genome. Run 2 was dedicated solely to GLST, i.e., no non-GLST libraries were simultaneously sequenced on the flow cell. The plot shows that run 1 and run 2 read-depths at each GLST base position (purple points) are highly correlated (Pearson’s r = 0.93, p < 0.001), and that run 1 had higher sequencing output than run 2. Read-depth values are square-root transformed and represent control sample TBM_2975_CL2_rep1.

**Supplementary Table 1.**
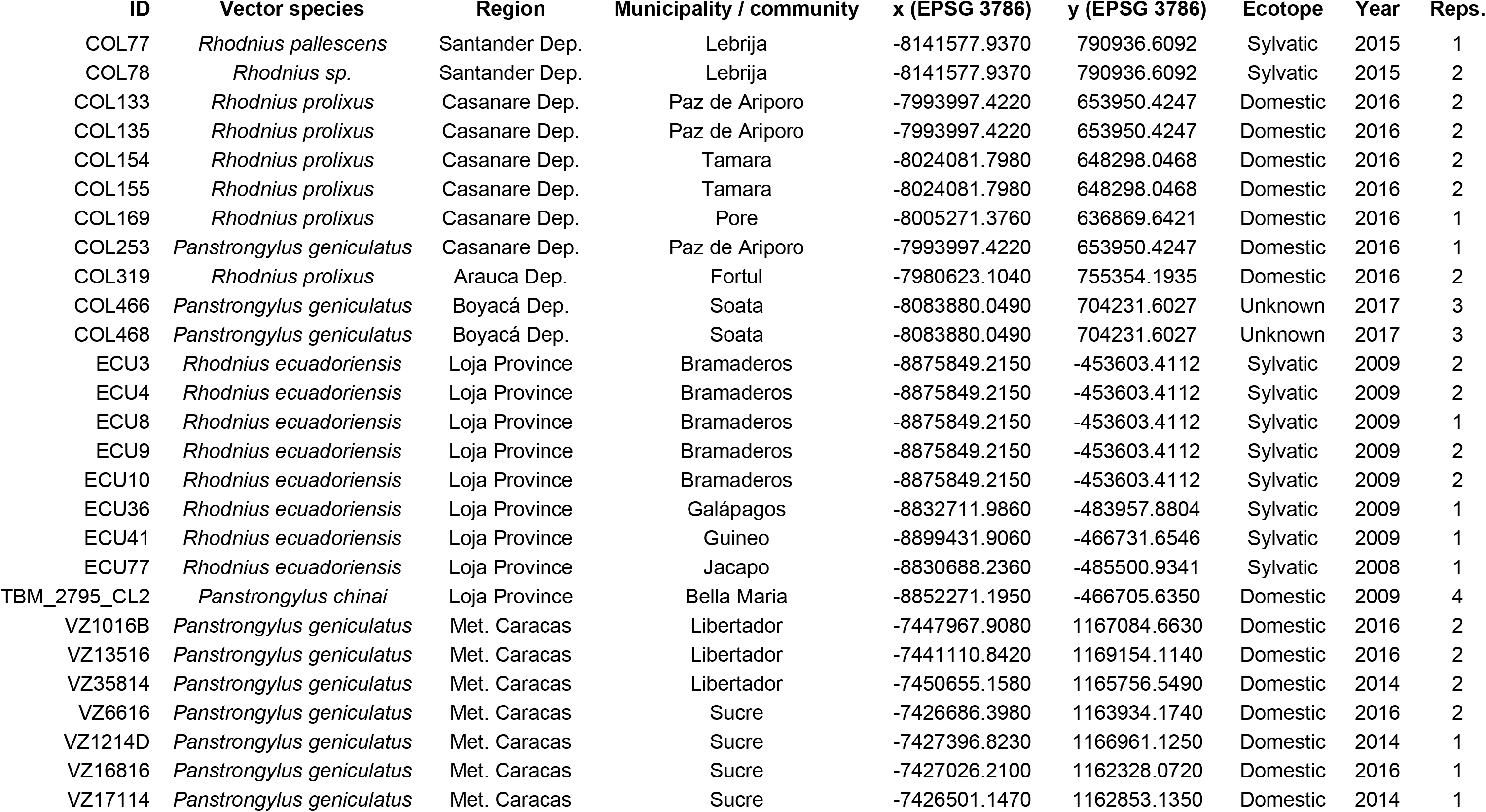
Details on *T. cruzi*-infected metagenomic triatomine gut samples from Colombia (COL), Venezuela (VZ) and Ecuador (ECU). Abbreviations: Dep. (Department); Met. Caracas (Metropolitan District of Caracas); EPSG (European Petroleum Survey Group coordinate system); reps. (technical replicates).

| ID | Vector species | Region | Municipality / community | x (EPSG 3786) | y (EPSG 3786) | Ecotope | Year | Reps. |
| --- | --- | --- | --- | --- | --- | --- | --- | --- |
| COL77 | *Rhodnius pallescens* | Santander Dep. | Lebrija | -8141577.9370 | 790936.6092 | Sylvatic | 2015 | 1 |
| COL78 | *Rhodnius sp.* | Santander Dep. | Lebrija | -8141577.9370 | 790936.6092 | Sylvatic | 2015 | 2 |
| COL133 | *Rhodnius prolixus* | Casanare Dep. | Paz de Ariporo | -7993997.4220 | 653950.4247 | Domestic | 2016 | 2 |
| COL135 | *Rhodnius prolixus* | Casanare Dep. | Paz de Ariporo | -7993997.4220 | 653950.4247 | Domestic | 2016 | 2 |
| COL154 | *Rhodnius prolixus* | Casanare Dep. | Tamara | -8024081.7980 | 648298.0468 | Domestic | 2016 | 2 |
| COL155 | *Rhodnius prolixus* | Casanare Dep. | Tamara | -8024081.7980 | 648298.0468 | Domestic | 2016 | 2 |
| COL169 | *Rhodnius prolixus* | Casanare Dep. | Pore | -8005271.3760 | 636869.6421 | Domestic | 2016 | 1 |
| COL253 | *Panstrongylus geniculatus* | Casanare Dep. | Paz de Ariporo | -7993997.4220 | 653950.4247 | Domestic | 2016 | 1 |
| COL319 | *Rhodnius prolixus* | Arauca Dep. | Fortul | -7980623.1040 | 755354.1935 | Domestic | 2016 | 2 |
| COL466 | *Panstrongylus geniculatus* | Boyacá Dep. | Soata | -8083880.0490 | 704231.6027 | Unknown | 2017 | 3 |
| COL468 | *Panstrongylus geniculatus* | Boyacá Dep. | Soata | -8083880.0490 | 704231.6027 | Unknown | 2017 | 3 |
| ECU3 | *Rhodnius ecuadoriensis* | Loja Province | Bramaderos | -8875849.2150 | -453603.4112 | Sylvatic | 2009 | 2 |
| ECU4 | *Rhodnius ecuadoriensis* | Loja Province | Bramaderos | -8875849.2150 | -453603.4112 | Sylvatic | 2009 | 2 |
| ECU8 | *Rhodnius ecuadoriensis* | Loja Province | Bramaderos | -8875849.2150 | -453603.4112 | Sylvatic | 2009 | 1 |
| ECU9 | *Rhodnius ecuadoriensis* | Loja Province | Bramaderos | -8875849.2150 | -453603.4112 | Sylvatic | 2009 | 2 |
| ECU10 | *Rhodnius ecuadoriensis* | Loja Province | Bramaderos | -8875849.2150 | -453603.4112 | Sylvatic | 2009 | 2 |
| ECU36 | *Rhodnius ecuadoriensis* | Loja Province | Galápagos | -8832711.9860 | -483957.8804 | Sylvatic | 2009 | 1 |
| ECU41 | *Rhodnius ecuadoriensis* | Loja Province | Guineo | -8899431.9060 | -466731.6546 | Sylvatic | 2009 | 1 |
| ECU77 | *Rhodnius ecuadoriensis* | Loja Province | Jacapo | -8830688.2360 | -485500.9341 | Sylvatic | 2008 | 1 |
| TBM_2795_CL2 | *Panstrongylus chinai* | Loja Province | Bella Maria | -8852271.1950 | -466705.6350 | Domestic | 2009 | 4 |
| VZ1016B | *Panstrongylus geniculatus* | Met. Caracas | Libertador | -7447967.9080 | 1167084.6630 | Domestic | 2016 | 2 |
| VZ13516 | *Panstrongylus geniculatus* | Met. Caracas | Libertador | -7441110.8420 | 1169154.1140 | Domestic | 2016 | 2 |
| VZ35814 | *Panstrongylus geniculatus* | Met. Caracas | Libertador | -7450655.1580 | 1165756.5490 | Domestic | 2014 | 2 |
| VZ6616 | *Panstrongylus geniculatus* | Met. Caracas | Sucre | -7426686.3980 | 1163934.1740 | Domestic | 2016 | 2 |
| VZ1214D | *Panstrongylus geniculatus* | Met. Caracas | Sucre | -7427396.8230 | 1166961.1250 | Domestic | 2014 | 1 |
| VZ16816 | *Panstrongylus geniculatus* | Met. Caracas | Sucre | -7427026.2100 | 1162328.0720 | Domestic | 2016 | 1 |
| VZ17114 | *Panstrongylus geniculatus* | Met. Caracas | Sucre | -7426501.1470 | 1162853.1350 | Domestic | 2014 | 1 |

**Supplementary Table 2.**
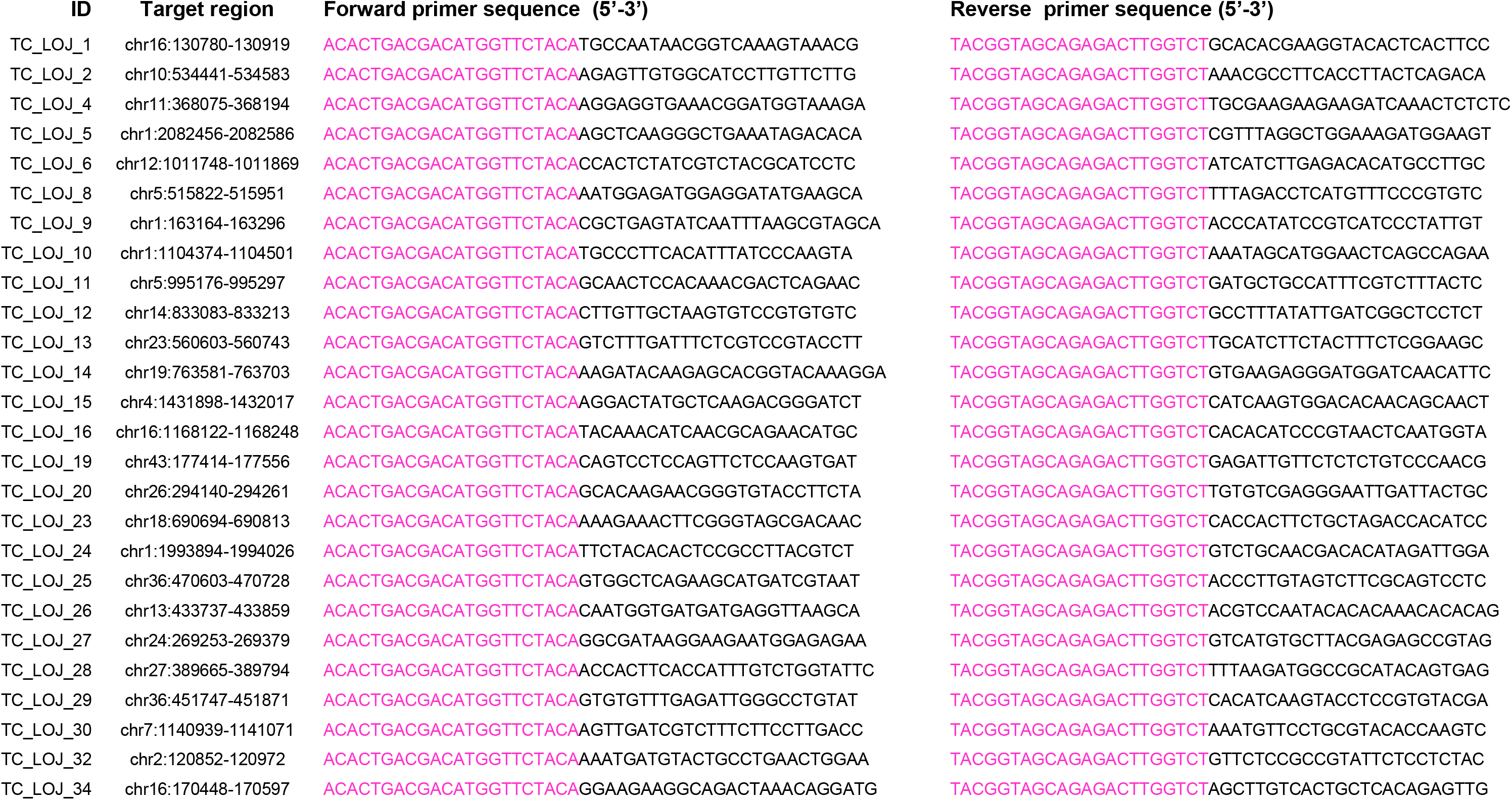
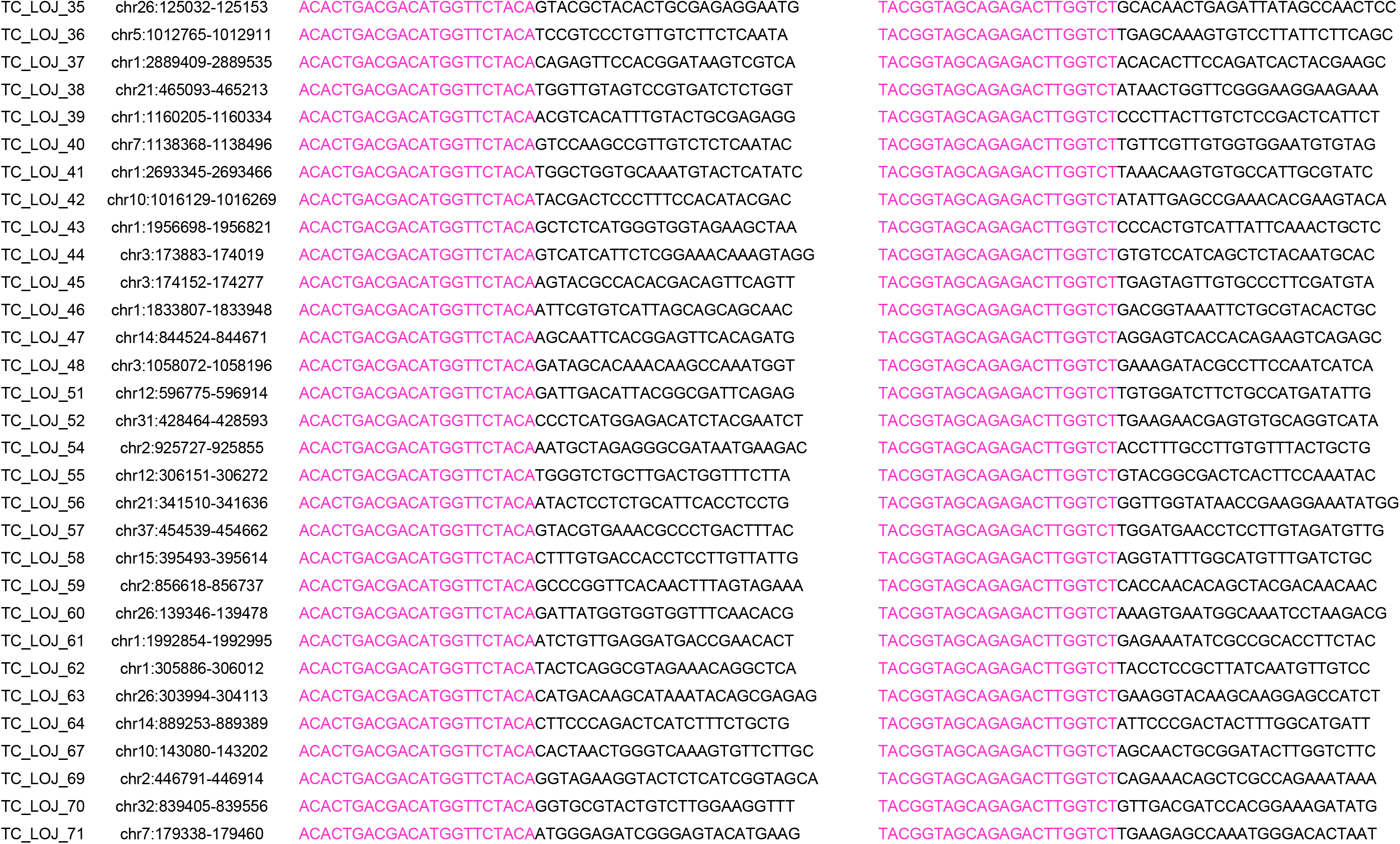
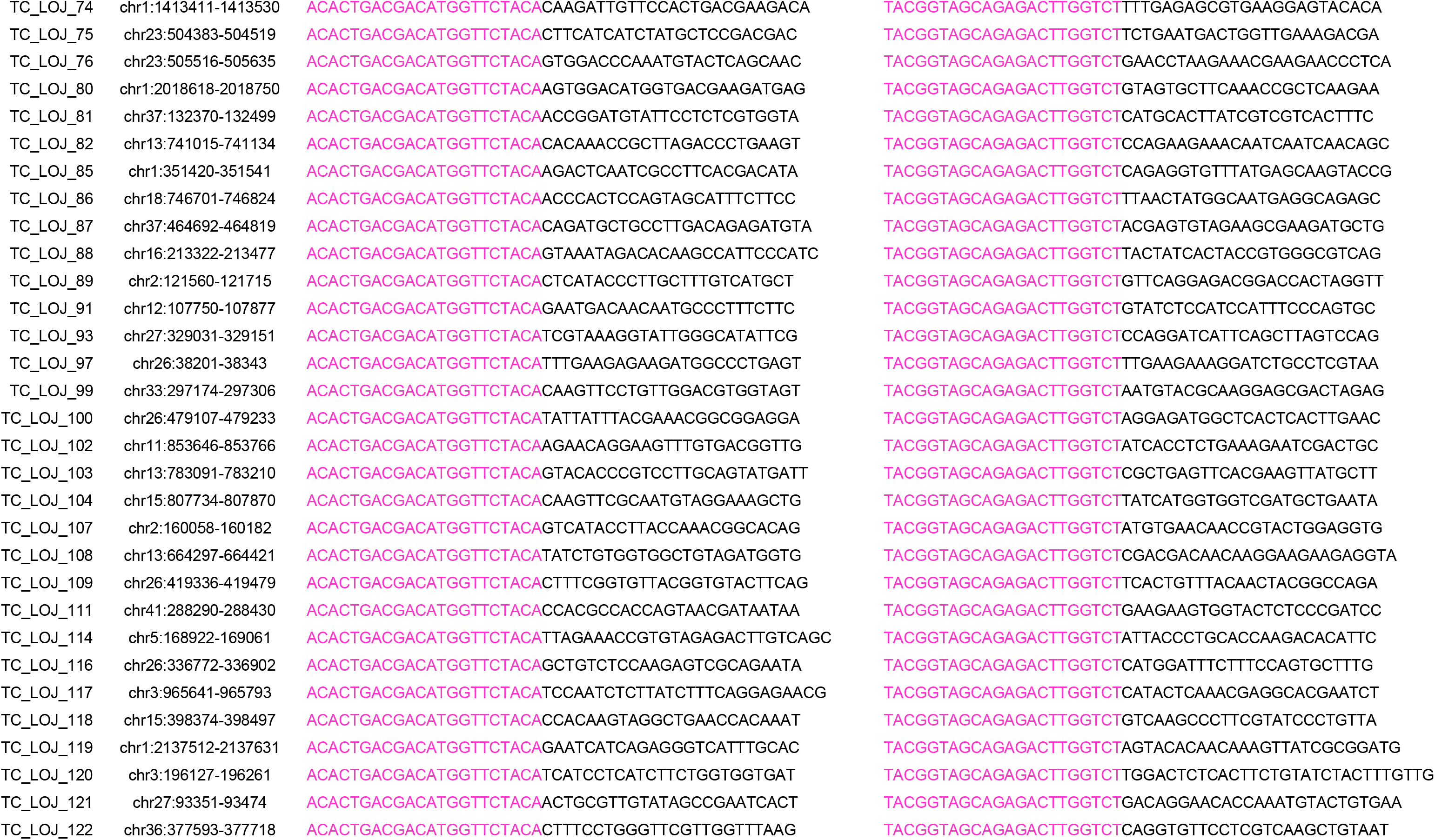
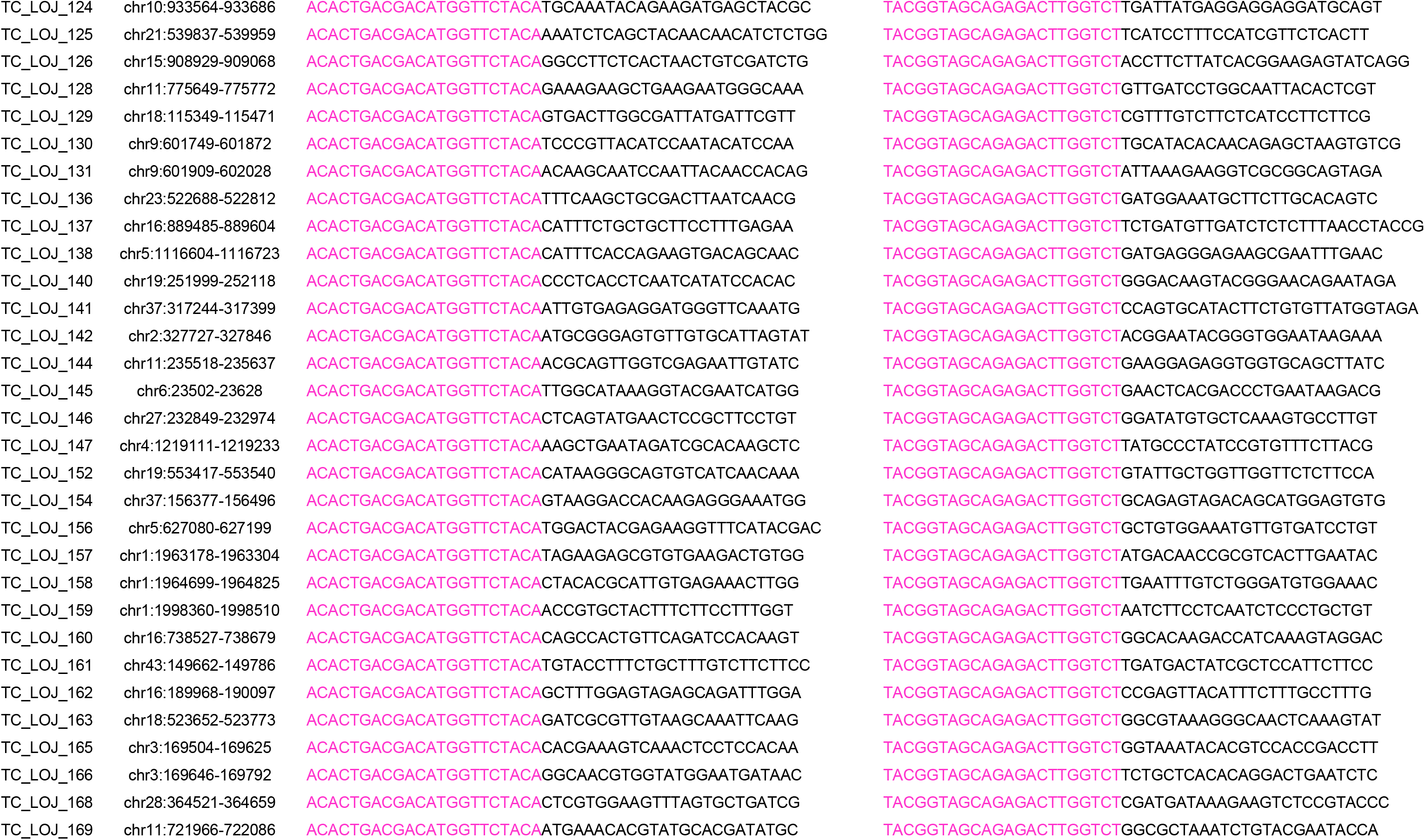
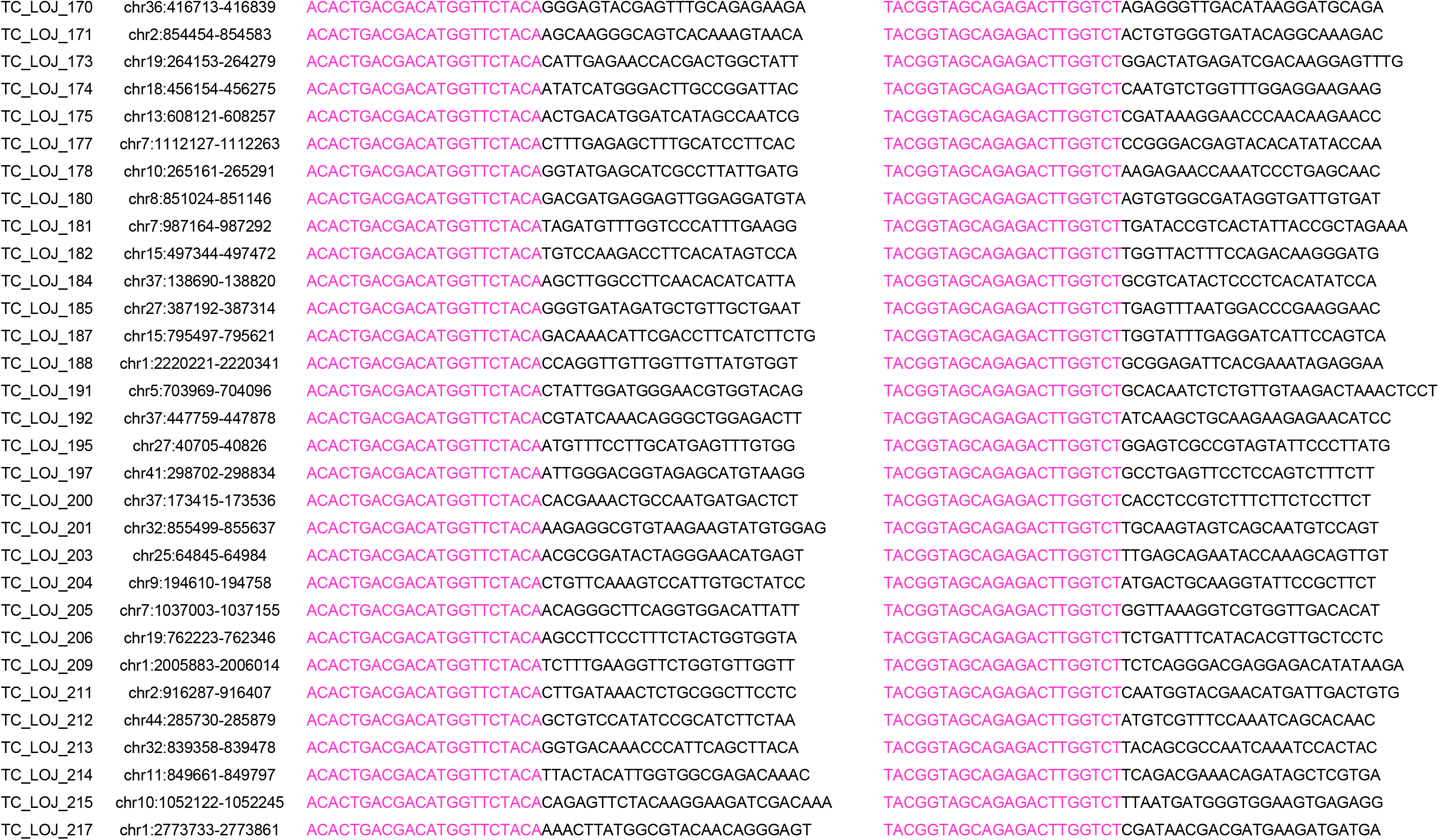
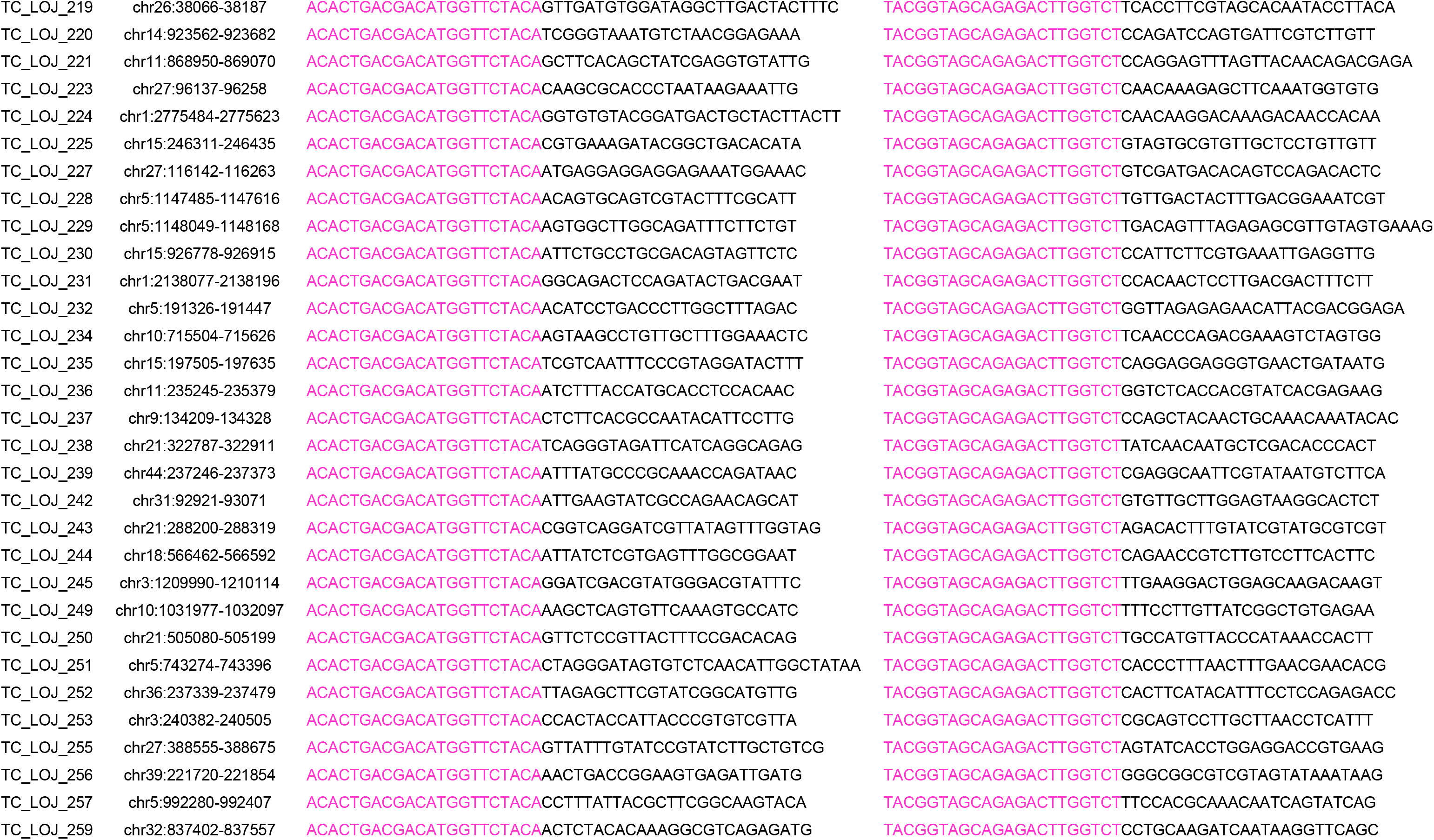
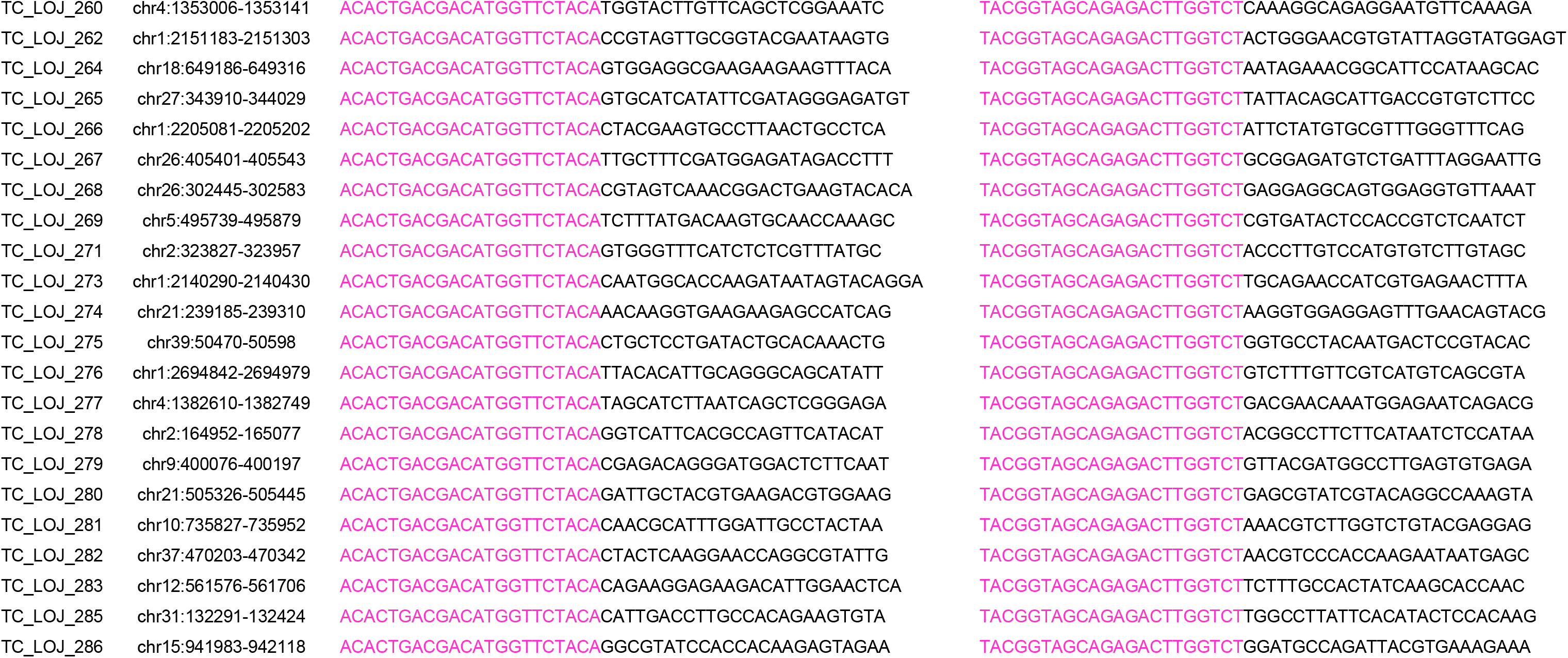
GLST primer sequences. The 3’ end of each first-round PCR primer is target-specific. The 5’ end of each forward primer contains CS1. The 5’ end of each reverse primer contains CS2. These sequencing primer binding sites are shown in pink. In subsequent barcoding PCR, the reverse primer consists of 5’-CAAGCAGAAGACGGCATACGAGAT*X*TACGGTAGCAGAGACTTGGTCT-3’, where *X* is a unique 10 nt barcode used to label each sample’s sequence reads. The reverse barcoding primer also contains CS2. The forward barcoding primer (5’-AATGATACGGCGACCACCGAGATCTACACTGACGACATGGTTCTA-3’) contains CS1 and is the same for all samples.

**Supplementary Table 3.** Summary of GLST library preparation and sequencing costs. Green dots indicate items/costs related to first-round PCR and clean-up. Blue dots indicate items/costs related to barcoding PCR and clean-up. The cost summary does not consider qPCR materials because we applied qPCR only for purposes of method development. It is not essential for GLST. Abbreviations: EUG Eurofins Genomics); NEB (New England Biolabs); MGRD (median genotype read-depth).

|  | Item | Availability<br>(quantity / price) | Quantity for<br>100 samples | Cost for<br>100 samples | Comment |
| --- | --- | --- | --- | --- | --- |
| Library preparation | 200 GLST primer primer pairs (EUG) ● | 60.90 ml / 1508.88 £ | 25 pmol | 1.26 £ | 18,861 bases purchased salt-free at 0.08 £ / base; primers delivered at 200 µM in 150 µl |
|  | Q5 High-Fidelity 2X Master Mix (NEB) ● | 2.5 ml / 106.75 £ | 500 µl | 21.35 £ |  |
|  | UltraPure Agarose (Invitrogen) ● | 100 g / 124.00 £ | 15.6 g | 19.34 £ | 13 agarose gels (0.8%) to visualize 100 samples, separated by empty lanes |
|  | 100 bp DNA Ladder* (NEB) ● | 50 ug / 34.50 £ | 13 ug | 8.97 £ | 0.5 ug ladder at left and right margins of each gel |
|  | 6X Gel Loading Dye (NEB) ● | 1 ml comes free with ladder* | 226 µl | 0.00 £ | 2 µl dye for each sample/ladder lane |
|  | PureLink Quick Gel Extraction Kit (Invitrogen) ● | 3 x 50 units / 143.64 £ | 100 units | 95.76 £ |  |
|  | SYBR Safe (Invitrogen) ● | 400 µl / 62.78 £ | 60 µl | 9.42 £ |  |
|  | Miscellaneous ● | n/a | n/a | 50.00 £ | Pipette tips, vials, blades, etc. |
|  | Barcoded reverse primer (EUG) ● | 0.02 µmol / 49.95 £ | 0.8 nmol | 2.00 £ | Primers purified by manufacturer using high performance liquid chromatography |
|  | Universal forward primer (EUG) ● | 0.02 µmol / 49.95 £ | 0.8 nmol | 2.00 £ | Primers purified by manufacturer using high performance liquid chromatography |
|  | Q5 High-Fidelity 2X Master Mix (NEB) ● | (see above) | 1 ml | 42.70 £ |  |
|  | Nuclease-free dH<sub>2</sub>O (Qiagen) ● | 1000 ml / 35.68 £ | 540 µl | 19.27 £ |  |
|  | Qubit assay tubes (Invitrogen) ● | 500 tubes / 51.50 £ | 102 tubes | 10.51 £ |  |
|  | Qubit dsDNA HS Assay Kit (Invitrogen) ● | 100 assay kit / 66.25 £ | 100 assays | 66.25 £ |  |
|  | UltraPure Agarose (Invitrogen) ● | (see above) | 1.2 g | 1.49 £ | Only one agarose gel (0.8%) is needed because samples have been pooled |
|  | 100 bp DNA Ladder (NEB) ● | (see above) | 1 ug | 0.69 £ | 0.5 ug ladder at left and right margins of the gel |
|  | 6X Gel Loading Dye (NEB) ● | (see above) | 9 µl | 0.00 £ | 7 µl dye for sample (pool) lane, 2 µl for each ladder lane |
|  | PureLink Quick Gel Extraction Kit (Invitrogen) ● | (see above) | 1 unit | 0.96 £ | Only one unit is needed because samples have been pooled |
|  | SYBR Safe (Invitrogen) ● | (see above) | 10 µl | 1.57 £ |  |
|  | Miscellaneous ● | n/a | n/a | 50.00 £ | Pipette tips, vials, blades, etc. |
| | Total library preparation cost for 100 samples: 256.41 £ | | | | ~ 3.15 \$ per sample |
| Sequencing | Item | Availability<br>(quantity / price) | Quantity for<br>100 samples | Cost for<br>100 samples | Comment |
|  | Illumina Reagent Kit v2 Micro | 1 cartridge / 390.00 £ | 1 cartridge | 390.00 £ | As listed at https://emea.illumina.com (March 2020) |
|  | 300-cycle Illumina MiSeq | 1 run / 40.00 £ | 1 run | 400.00 £ | Costs for quality control, data storage, etc. vary considerably among providers |
| | Total sequencing cost for 100 samples: 790.00 £ | | | | ~ 9.72 \$ per sample; 70x MGRD expected based on 125x MGRD for 56 samples in run 2 |

